# Catalase 2-dependent regulation of autophagy in response to carbon starvation in Arabidopsis

**DOI:** 10.64898/2026.04.15.718673

**Authors:** Chi Zhang, Song-Qi Li, Pengwei Jing, Ji-Xiao Wu, Bo-Hao Chen, Cai-Yi Liao, Hong Yang, Kai-Kai Lu, Ru-Feng Song, Christine H. Foyer, Wen-Cheng Liu

**Affiliations:** State Key Laboratory of Crop Stress Adaptation and Improvement, Collaborative Innovation Center of Crop Stress Biology, College of Life Sciences, Henan University, Kaifeng 475004, China; School of Biosciences, College of Life and Environmental Sciences, University of Birmingham, Edgbaston B15 2TT, UK

## Abstract

Reactive oxygen species (ROS) are fundamental regulators of plant development and stress acclimation. However, their roles in carbon starvation tolerance remain largely unresolved. We therefore dissected the dynamics of redox regulation in carbon-starved Arabidopsis leaves. We present data showing the presence of an early ROS wave that activates autophagy, followed by a sustained, accumulation of hydrogen peroxide (H_2_O_2_) that leads to programmed cell death. We show that catalase 2 (CAT2) constitutively binds and sequesters the autophagy initiators ATG1a and ATG13a, a process that prevents the activation of autophagy under carbon-replete conditions. The accumulation of H_2_O_2_ in response to carbon starvation leads to oxidation of the Cys-370 and Cys-413 residues on the CAT2 protein and dissociation from ATG1a/13a, thereby allowing ATG1-kinase complex assembly and autophagosome formation. The expression of CAT2-C370S or CAT2-C413S variants in the *cat2-1* mutant significantly enhanced carbon starvation tolerance, because both variants retain catalase activity but fail to bind ATG1a/13a. We conclude that CAT2 functions as a redox-sensitive regulator of autophagy.

## Introduction

Autophagy is a conserved pathway in eukaryotic organisms that degrades excess cellular components and/or damaged organelles. It is dependent on vesicular transport and vacuolar enzymes, facilitating recycling of resources and maintaining cellular homeostasis (Yoshimoto et al., 2014). Autophagy is crucial to plant biology, fulfilling key roles in plant growth, development, and stress responses (Yang et al., 2020). At least three autophagy pathways have been identified in plants: macroautophagy, selective autophagy, and microautophagy (Zhang et al., 2026). Of these, macroautophagy is the predominant process in plant cells. The macroautophagy pathway starts with the initiation and nucleation of phagosomes, which engulf degradation cargo and extend membranes to form autophagosomes with characteristic double-membrane structures (Zhang et al., 2026). Subsequently, the autophagosome fuses with the vacuole via the vesicular transport system, ultimately releasing its contents into the vacuolar lumen for degradation (Yoshimoto and Ohsumi, 2018). Phagophore formation is instigated by the formation of functional protein complexes composed of core autophagy proteins, including: (1) the ATG1-ATG13 protein kinase complex; (2) the ATG6-phosphatidylinositol 3-kinase (PI3K) complex; (3) the ATG9 membrane delivery complex; and (4) two ubiquitin-like conjugation systems (ATG5-ATG12 and ATG8-phosphatidylethanolamine) (Yoshimoto and Ohsumi, 2018). Mutants in autophagy genes exhibit phenotypes, such as shortened growth periods, leaf chlorosis, male sterility and stunted plant growth, particularly under stress conditions (Kurusu et al., 2014; Minina et al., 2018; Wada et al., 2015).

The ATG1-13 kinase complex is responsible for the initiation of autophagy, which is regulated by a series of post-translational modification (PTM) switches. Under nutrient-sufficient conditions, ATG13 is phosphorylated by the target protein of rapamycin (TOR) kinase (Wang et al., 2022), which causes dissociation of the ATG1 and ATG13 proteins, preventing the assembly of the ATG1-ATG13 protein complex and thereby inhibiting autophagy (Suttangkakul et al., 2011). Under nutrient stress, the energy-sensing protein SnRK1, which is upstream of TOR, detects changes in cellular energy status and inactivates the TOR kinase (Nukarinen et al., 2016; Otegui et al., 2017). The resultant dephosphorylation of ATG13 promotes phosphorylation of ATG1. Recruitment of the scaffold proteins ATG11 and ATG101 allows the formation of the mature ATG1-13 protein complex, initiating autophagy (Suttangkakul et al., 2011; Wang and Hou, 2022). Previous studies have shown that ATG13 is a regulatory subunit in this complex that undergoes both phosphorylation by the TYPE ONE PROTEIN PHOSPHATASE (TOPP) and ubiquitination by SINAT1 and SINAT2 (Qi et al., 2022; Qi et al., 2017; Wang et al., 2022). These two processes act in opposition regulating the stability and activation of the ATG1-ATG13 kinase complex, which drives autophagosome formation.

Reactive oxygen species (ROS) such as H_2_O_2_ are important redox signals that regulate autophagy by oxidizing protein cysteine (Cys) residues and redox posttranslational modifications (Mittler et al., 2011). In animals, hydrogen peroxide (H_2_O_2_) enhances autophagy by oxidizing HsATG4b at Cys-78 site, thereby inhibiting ATG4 dissociation from ATG8-PE (Scherz-Shouval et al., 2007). A similar mechanism also occurs in yeast and Chlamydomonas, (Perez-Perez et al., 2016; Perez-Perez et al., 2014). Such findings demonstrate that ATG4 activity can be regulated by H_2_O_2_-mediated oxidation of specific cysteine residues. The autophagy pathway in higher plants is also regulated by H_2_O_2_, although the precise mechanisms that are involved have not been demonstrated (Woo et al., 2014). There is a close regulatory relationship between autophagy and ROS accumulation in plants (Zhang et al., 2026). In Arabidopsis, the autophagy mutants *atg5* and *atg7* show high levels of oxidation under short-day conditions (Yoshimoto et al., 2009). Similarly, *RNAi-ATG6* barley plants showed significant nutritional stress and enhanced sensitivity to oxidation (Zeng et al., 2017), indicating reciprocal control between the autophagy pathway and oxidative metabolism. Autophagy levels were increased in Arabidopsis following treatment with the prooxidant methyl viologen (MV) or with H_2_O_2_ (Xiong et al., 2007).

Catalase 2 (CAT2) is the most important catalase isoform in leaves (Baker et al., 2023), catalyzing the breakdown of H_2_O_2_ produced by photorespiration and other oxidative pathways into water and oxygen (Wang et al., 2024). As such, CAT2 is a key antioxidant enzyme, plants exposed to abiotic stresses, such as salinity, drought, and high light (Fu et al., 2023; Michelet et al., 2013; Song et al., 2021; Yang et al., 2025). Our previous studies have shown that CAT2 enhances the activity of both acyl-CoA oxidase (ACX) 2 and ACX3 through direct interactions, that regulate seedling growth and immune responses (Liu et al., 2017; Yuan et al., 2017; Zhang et al., 2021). Previous studies have also shown that CAT2 is able to associate with many proteins in the cytosol (Baker et al., 2023), and that it can be relocated to the nucleus (Al-Hajaya et al., 2022). Furthermore, CAT2 forms biomolecular condensates with other cytosolic proteins in a redox-dependent manner, in a way that upregulates CAT activity, and regulates CAT2 transport to the nucleus (Lin et al., 2025).

In the present study, we have investigated the role of CAT2 and of H_2_O_2_ in the regulation of autophagy triggered by carbon starvation in Arabidopsis. We show that carbon deficiency results in H_2_O_2_ accumulation at an early stage and that this stimulates autophagy. In contrast, further increases in H_2_O_2_ levels at the later stages of carbon starvation leading to chlorosis. We show that CAT2 blocks the assembly of the ATG1-ATG13 complex through redox-dependent protein/protein interactions in the absence of stress. Carbon starvation triggers H_2_O_2_ accumulation, which disrupts CAT2 binding to ATG1 and ATG13 through oxidation of the Cys-370 and Cys-413 on the CAT2 protein, thereby promoting autophagy. Taken together, these findings demonstrate that the redox-dependent binding of CAT2 to ATG1 and ATG13 regulates autophagy, through a Cys-dependent redox switch mechanism.

## RESULTS

### H_2_O_2_ regulates the early and late stages of carbon starvation

To determine how carbon starvation affects H_2_O_2_ accumulation in Arabidopsis, seedlings were grown on sucrose-depleted MS medium in the dark. H_2_O_2_ accumulated in both the shoots and roots of the wild-type (WT) seedlings as a result of carbon deprivation, as indicated by 3,3′-diaminobenzidine (DAB) staining (Figure 1A-B). To identify the source of H_2_O_2_ production induced by carbon starvation, we compared H_2_O_2_ accumulation in the *cat2-1* mutants and *rbohd/f* mutants that are deficient in two of the plasmalemma-localized respiratory burst oxidase homologues (RBOH) (Song et al., 2021). The *cat2-1* mutants accumulated more H_2_O_2_ than the WT under conditions of carbon starvation treatment. In contrast, H_2_O_2_ accumulation was decreased relative to the WT in the *rbohd/f* mutants (Figure 1A-B). We next examined whether RBOH activity was changed in response to starvation using in-gel assays. RBOH activity increased at the early stages of carbon starvation and then declined gradually as the starvation period progressed (Figure S1A-B). RBOH activity was markedly decreased in the *rbohd/f* mutants (Figure S1C). In contrast, CAT activity increased within the first day of starvation, followed by a biphasic response (Figure S1D).

**Figure 1.**
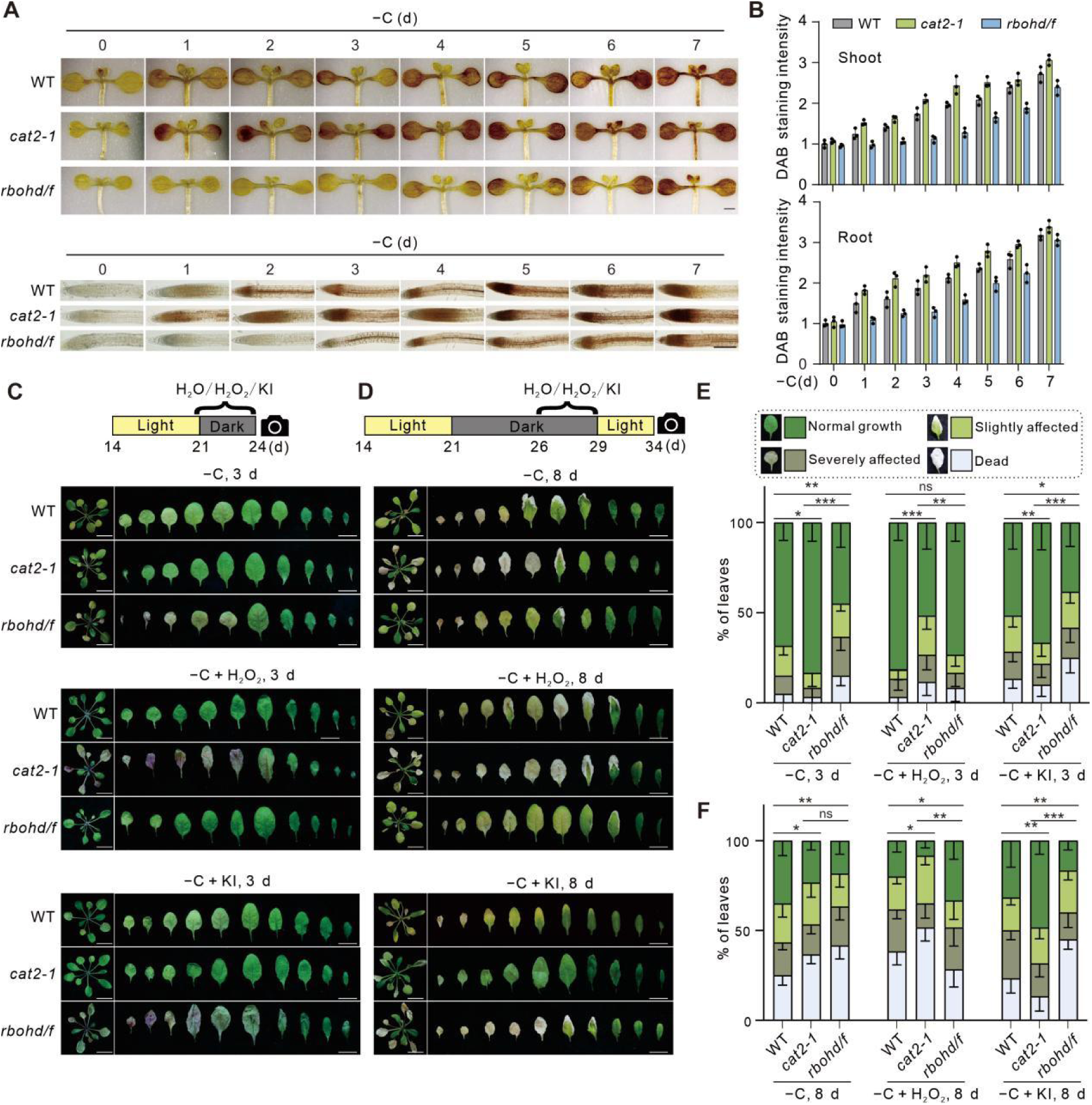
H_2_O_2_ displays a dual role in plant carbon starvation tolerance. (**A**) DAB staining images of WT, *cat2-1* and *rbohd/f* seedling shoots (above) and roots (below) during a time-course analysis of carbon starvation treatment (days indicated above). One-week-old WT, *cat2-1*, and *rbohd/f* mutant seedlings grown on sucrose-containing MS medium under long-day conditions were transferred to sucrose-lacking MS medium under dark (-C) for the indicated days, and subjected to DAB staining. Bar = 5 mm. (**B**) The relative DAB staining intensity in (A). The DAB staining intensity of the untreated wild-type (WT) leaves was set to 1. Data are means ± SD (n=3). (**C-D**) Representative images of the WT, *cat2-1* and *rbohd/f* plants treated with carbon starvation for 3 d (C) or 8 d (D) in the presence of H_2_O, H_2_O_2_ or KI. Bar = 1 cm. 21-d-old soil-grown WT, *cat2-1*, and *rbohd/f* plants grown under long-day conditions were transferred to dark conditions, and subsequently sprayed with H_2_O, 0.5 mM H_2_O_2_, or 0.5 mM KI. Treatments were applied either during days 21–24 (C) or days 26–29 (D). Bar = 1 cm. (**E-F**) Representative leaves used in carbon starvation phenotypic analysis, including normal growth, slightly affected, severely affected and dead leaves. Percentages of the four types of leaves in the WT, *cat2-1* and *rbohd/f* mutant plants treated with carbon starvation for 3 d (E) and 8 d (F). Data are means ± SD (n=3). Asterisks indicate statistically significant differences (*P<0.05, **P<0.01 and ***P<0.001, by ANOVA); “ns” indicates not significant. Bar = 10 mm.

To determine whether the observed changes of H_2_O_2_ accumulation influence carbon starvation responses, we compared carbon starvation-induced chlorosis in the WT, *cat2-1* and *rbohd/f* mutants (Figure 1C-D). The WT leaves exhibited partial chlorosis on the 3^rd^ day (designated as an early stage of carbon starvation) and severe chlorosis on the 8^th^ day (designated as a late stage of carbon starvation) (Figure 1C-G). The leaves of the *cat2-1* mutant showed less chlorosis than the WT at the early stage but more chlorosis at the late stage (Figure 1C-F), suggesting that catalase and/or H_2_O_2_ exerts different time-dependent controls on the carbon starvation response. The leaves of the *rbohd/f* mutants exhibited stronger chlorosis than the WT at the early stages of the response but only slightly more chlorosis than the WT in the late stages (Figure 1C-F).

To explore the role of H_2_O_2_ at different stages of the carbon starvation response further, we manipulated endogenous H_2_O_2_ levels by either adding H_2_O_2_ or the H_2_O_2_-scavenger potassium Iodide (KI). The exogenous application of a low dose of H_2_O_2_ (less than 1 mM, to avoid oxidative stress and damages) alleviated leaf chlorosis in the WT and partially rescued the more sensitive phenotype in the *rbohd/f* mutants at the early stages (Figure 1C, E); However, the addition of H_2_O_2_ accelerated leaf chlorosis in the *cat2-1* mutants (Figure 1C, E). In contrast, KI treatment increased chlorosis in both WT and the *rbohd/f* mutants but not in the *cat2-1* mutants at the early stage of carbon starvation (Figure 1C, E). During the late carbon starvation stages, the exogenous H_2_O_2_ application enhanced chlorosis in all genotypes (Figure 1D, 1F). While KI treatment had little effect on chlorosis in the WT and *rbohd/f* mutants, it largely prevented late-stage chlorosis in the *cat2-1* mutants (Figure 1D, 1F).

### H_2_O_2_ enhances carbon starvation tolerance by facilitating autophagy

To test whether H_2_O_2_ activates carbon starvation responses through the induction of autophagy, we subjected WT and two previously reported autophagy mutants, *atg1abc* and *atg13ab*, to carbon deprivation in the presence of H_2_O_2_ or KI. Both the *atg1abc* and *atg13ab* mutants displayed hypersensitive phenotypes compared to the WT, under carbon starvation conditions (Figure 2A, 2B). Treatment with H_2_O_2_ significantly increased the stability of chlorophyll in the WT in response to carbon starvation, whereas KI treatment increased the loss of chlorophyll (Figure 2A, 2B). Neither H_2_O_2_ nor KI had any effect on the carbon starvation hypersensitive phenotype observed in both the *atg1abc* and *atg13ab* mutants (Figure 2A, 2B). This finding suggests that H_2_O_2_ promotes carbon starvation responses through the ATG1-and ATG13-mediated autophagy pathway. To test this hypothesis, we investigated the effects of H_2_O_2_ and KI on autophagosome formation in transgenic *35S::GFP-ATG8e* plants (Qi et al., 2022). The number of carbon starvation-induced autophagosomes was enhanced in the presence of H_2_O_2_ but suppressed by KI (Figure 2C, 2D). The flux through the autophagy pathway was enhanced in the presence of H_2_O_2_ but inhibited in the presence of KI under carbon deprivation conditions, as evidenced by a higher ratio of free GFP to GFP-ATG8e in H_2_O_2_-treated *35S::GFP-ATG8e* plants and a lower ratio in KI-treated plants, relative to the untreated controls (Figure 2E).

**Figure 2.**
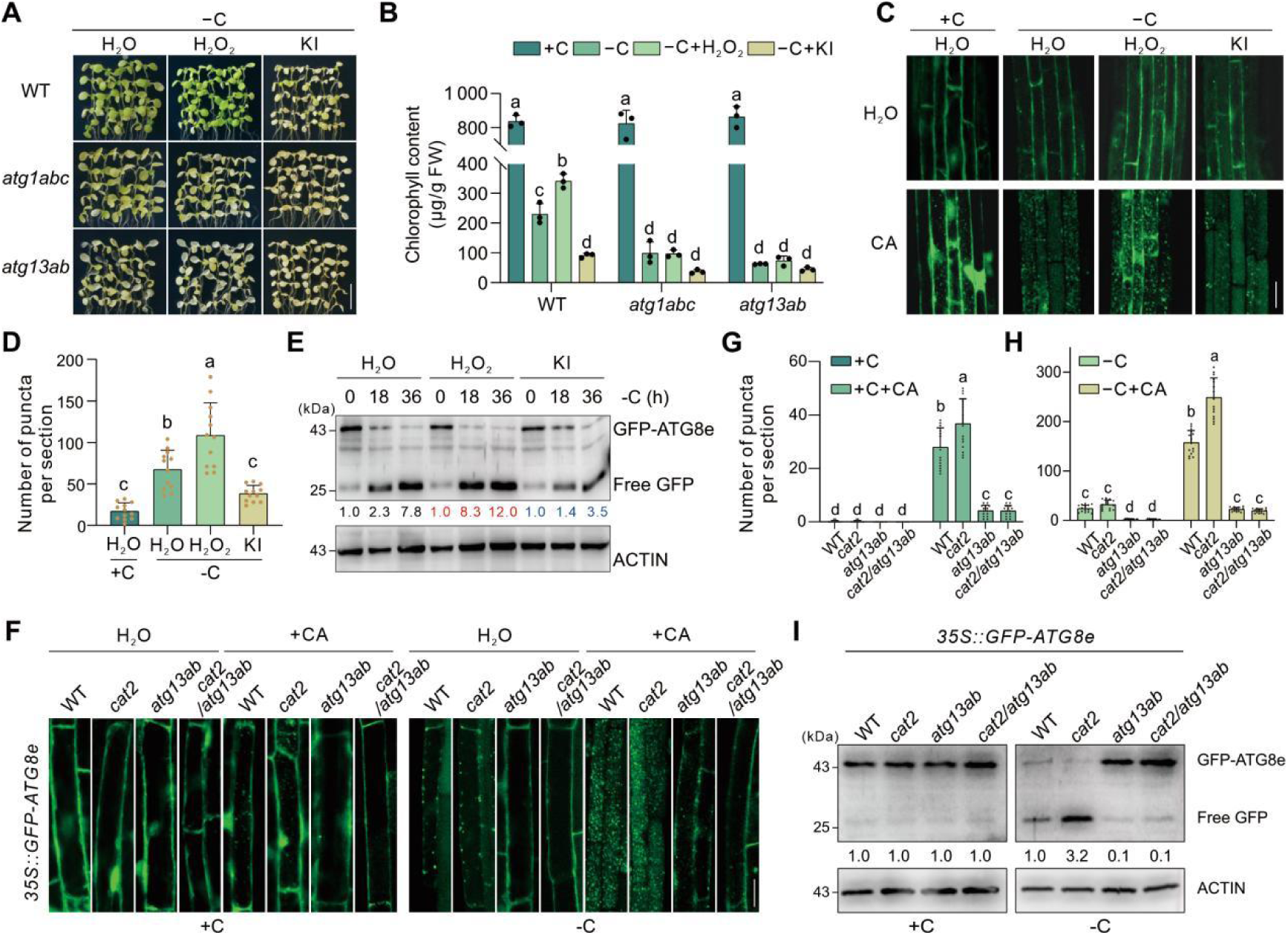
H_2_O_2_ promotes autophagic flux and enhances carbon starvation tolerance. (**A-B**) Phenotypes of the carbon-starved WT, *atg1abc*, and *atg13ab* mutant plants in the presence of H_2_O_2_ or KI. 7-d-old WT, *atg1abc*, and *atg13ab* mutant seedlings grown on sucrose-containing MS medium under long-day conditions were transferred to sucrose-lacking MS medium supplemented with mock (H_2_O), 0.1 mM H_2_O_2_ or 0.5 mM KI under dark for additional 5-7 d. Bar = 10 mm. Chlorophyll contents of the plants shown in (A) were analysed. Data are means ± SD calculated from 3 independent experiments. For each experiment, five entire plants (technical replicates) were used per genotype. Bars with different letters indicate significant differences at p < 0.05, revealed using a one-way analysis of variance with a Tukey’s multiple comparison test. (**C-D**) Confocal imaging analysis of the *35S::GFP-ATG8e* seedlings treated with +C, -C+H_2_O, -C+H_2_O_2_, or -C+KI. One-week-old *35S::GFP-ATG8e* seedlings grown on sucrose-containing MS medium under long-day conditions were subjected to sucrose-lacking MS medium supplemented with H_2_O (-C+H_2_O), 0.1 mM H_2_O_2_ (-C+H_2_O_2_) or 0.5 mM KI (-C+KI) in the presence or absence of 1 µM CA under dark, respectively for 16 h. Bar = 25 mm. The number of GFP puncta per root section in root cells was analysed (D). Data are means ± SD (n=12). Bars with different letters indicate significant differences at p < 0.05, revealed using a one-way analysis of variance with a Tukey’s multiple comparison test. (**E**) Immunoblot analysis showing the effect of H_2_O_2_ or KI on the processing of GFP-ATG8e of *35S::GFP-ATG8e* seedlings. One-week-old *35S::GFP-ATG8e* seedlings were treated with carbon starvation in the presence of 0.1 mM H_2_O_2_ or 0.5 mM KI for 0, 18, and 36 h. Total proteins of the seedlings were extracted and anti-GFP antibody was used for the immunoblot analysis. The position of GFP-ATG8e fusion and free GFP are indicated. The ratio between free GFP and GFP-ATG8e is shown below. Actin was used as the loading control. (**F-H**) Confocal imaging analysis showing the autophagosomes in the *35S::GFP-ATG8e*, *35S::GFP-ATG8e/cat2* and *35S::GFP-ATG8e/cat2/atg13ab* lines. 7-d-old *35S::GFP-ATG8e*, *35S::GFP-ATG8e/cat2* and *35S::GFP-ATG8e/cat2/atg13ab* seedlings grown on sucrose-containing MS medium under long-day conditions were subjected to carbon starvation (–C) conditions in the presence or absence of 1 µM CA for 16 h. The formation of autophagosomes was visualized by fluorescence confocal microscopy. The number of GFP puncta per section in root cells was analysed (G-H). (**I**) Immunoblot analysis of GFP–ATG8e and free GFP in *35S::GFP-ATG8e*, *35S::GFP-ATG8e/cat2* and *35S::GFP-ATG8e/cat2/atg13ab* lines under +C and −C conditions, respectively.The ratio between free GFP and GFP-ATG8e is shown below. Actin was used as the loading control.

We next evaluated the contribution of endogenous H_2_O_2_ to autophagic activity using the *cat2-1* mutants. We generated *35S::GFP-ATG8e cat2-1* plants by crossing the *35S::GFP-ATG8e* plants with the *cat2-1* mutants. The *35S::GFP-ATG8e cat2-1* plants had more autophagosomes and a higher autophagic flux under conditions of carbon starvation than the *35S::GFP-ATG8e* plants (Figure 2F–2H). We also generated *35S::GFP-ATG8e atg13ab* and *35S::GFP-ATG8e atg13ab cat2-1* plants through genetic crosses and subjected them to carbon starvation treatment. The carbon starvation-induced increase in both autophagosomes and autophagic flux were largely blocked in both the *35S::GFP-ATG8e atg13ab* and *35S::GFP-ATG8e atg13ab cat2-1* plants compared to the *35S::GFP-ATG8e* and *35S::GFP-ATG8e cat2-1* plants (Figure 2F–2I). This finding suggests that the enhanced autophagic activity observed in *cat*2*-1* mutants depends on ATG13.

### CAT2 interacts with both ATG1 and ATG13 to inhibit carbon starvation tolerance

To determine the role of CAT2 in the regulation of autophagy, we used CAT2 as a bait to screen for interacting proteins in a yeast-two hybrid system. This screen identified ATG13a (Figure 3A), a protein that functions as a component of the autophagy initiation complex through association with ATG1. CAT2 also interacted with ATG13b and the three members of ATG1 family, including ATG1a, ATG1b, and ATG1c (Figure 3A, S2E). To further verify these interactions, bimolecular fluorescence complementation (BiFC) assays were performed, in which CAT2 fused to the N-terminal half of yellow fluorescent protein (nYFP), was co-expressed with ATG1a or ATG13a fused to the C-terminal half of YFP (cYFP) in *N. benthamiana* leaves. Reconstituted YFP fluorescence was observed in the leaves co-expressing nYFP-CAT2 and cYFP-ATG1a or cYFP-ATG13a, but there was no detectable fluorescence in the negative controls (Figure 3B). The interaction between CAT2 and ATG1a *in planta* was also verified by co-immunoprecipitation (Co-IP) assays in which CAT2 and GFP-tagged ATG1a were co-expressed in *N. benthamiana* leaves and then immunoprecipitated using an anti-GFP antibody. CAT2 was specifically detected by the anti-CAT2 antibody in the protein precipitates using the anti-GFP antibody (Figure 3C). To test these interactions *in vitro*, we purified glutathione S-transferase (GST)-tagged ATG1a and 6×His-tagged CAT2 proteins in *Escherichia coli* (*E. coli*) and we performed GST pull-down assays using glutathione-sepharose beads. The 6×His-CAT2 was specifically pulled down by GST-ATG1a, but not by the GST tag alone (Figure 3D). These results demonstrate that CAT2 interacts with ATG1a both *in vivo* and *in vitro* assays. The BiFC and GST pull-down assays also showed that CAT2 interacts with ATG13a *in vivo* and *in vitro* (Figure 3B, 3E). In addition, ATG1a interacts with ATG13a both *in vivo* and *in vitro* (Figure 3B, 3F), consistent with previous reports (Hama et al., 2025; Qi et al., 2022). These analyses suggest that CAT2, ATG1 and ATG13 may form a ternary protein complex.

**Figure 3.**
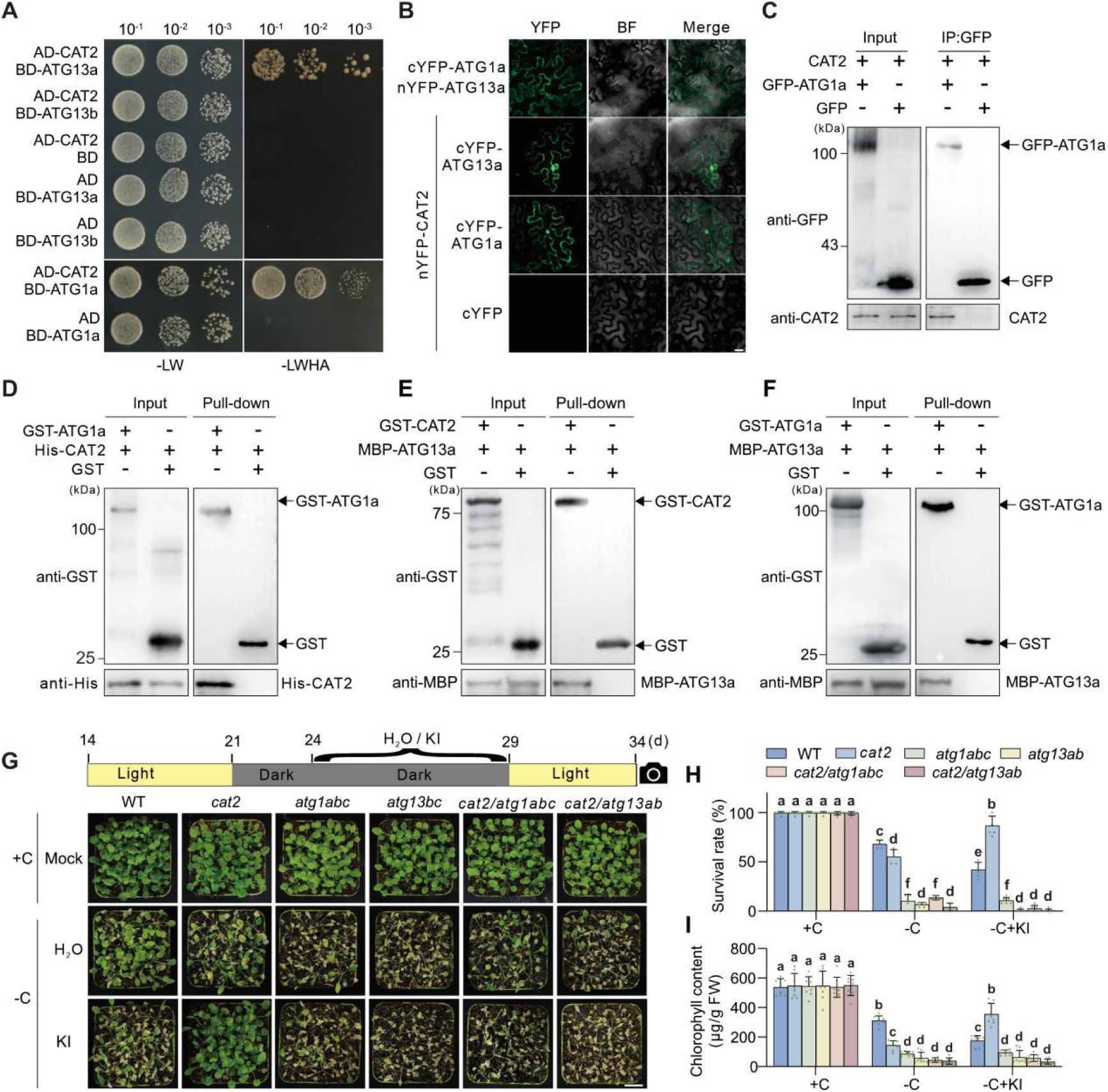
CAT2 physically interacts with ATG1a and ATG13a and genetically modulates autophagy-dependent tolerance. (**A**) Yeast two-hybrid assay showing the interaction between CAT2 and ATG1a or ATG13a. (**B**) BiFC assay showing the interaction among ATG1a, ATG13a, and CAT2. CAT2 fused to nYFP and ATG1a or ATG13a fused to cYFP were co-expressed in *N. benthamiana* leaves and the reconstituted YFP fluorescence was observed with confocal microscopy. Scale bars = 20 μm. (**C**) Co-immunoprecipitation (Co-IP) assay showing the interaction between CAT2 and ATG1a. GFP or GFP-ATG1a was co-expressed with CAT2-MYC in *N. benthamiana* leaves for 3 d. The total proteins were extracted and immunoprecipitated with anti-GFP antibody, and the precipitate was assayed with anti-GFP and anti-CAT2 antibodies, respectively. (**D**) GST pull-down assay showing the interaction between CAT2 and ATG1a. The purified 6×His–CAT2 was mixed with the purified GST–ATG1a or GST, and immobilized on glutathione sepharose beads on ice. After washing, the eluted proteins were subjected to immunoblot analysis with anti-GST or anti-His antibodies, respectively. (**E**) GST pull-down assay showing the interaction between GST–CAT2 and MBP-ATG13a. The purified GST or GST–CAT2 was mixed with the purified MBP–ATG13a, and immobilized on glutathione sepharose beads on ice. After washing, the eluted proteins were subjected to immunoblot analysis with anti-GST or anti-MBP antibodies, respectively. (**F**) GST pull-down assay showing the interaction between GST–ATG1a and MBP-ATG13a. The purified GST or GST–ATG1a was mixed with the purified MBP–ATG13a, and immobilized on glutathione sepharose beads on ice. After washing, the eluted proteins were subjected to immunoblot analysis with anti-GST or anti-MBP antibodies, respectively. (**G–I**) Phenotypes (**G**) of the WT*, cat2, atg1abc, atg13ab, cat2/atg1abc* and *cat2/atg13ab* mutant plants treatment of carbon starvation with H_2_O or KI. Three-week-old soil-grown WT*, cat2, atg1abc, atg13ab, cat2/atg1abc* and *cat2/atg13ab* mutant plants under long-day conditions were transferred to darkness and spayed with H_2_O or 0.5 mM KI as indicated in the flow chart. The treated plants were transferred to long-day conditions for recovery, and their survival rates (H) and chlorophyll content were analysed (I). Data are means ± SD (n = 5). Bars with different letters indicate significant differences at p < 0.05, revealed using a one-way analysis of variance with a Tukey’s multiple comparison test.

To confirm the relationships between *CAT2* and *ATG1a* or *ATG13a*, we generated *cat2-1 atg1abc* and *cat2-1 atg13ab* mutants via genetic crossing of *cat2-1* with *atg1abc* or *atg13ab*. All lines were then subjected to carbon starvation. Like the *atg1abc* and *atg13ab* mutants, the *cat2-1 atg1abc* and *cat2-1 atg13ab* mutants showed hypersensitivity to carbon starvation after a long period as evidenced by survival rates and chlorophyll contents (Figure 3G-3I). To exclude the extra effect of higher H_2_O_2_ and focus on how disruption of CAT2 protein affects autophagy in the *cat2-1* mutants, KI was employed to treat the WT, *cat2-1*, *atg1abc*, *atg13ab*, *cat2-1 atg1abc* and *cat2-1 atg13ab* mutants, reducing excess ROS in the *cat2-1* mutants. In the presence of KI, the *cat2-1* mutant had higher survival rate and chlorophyll content than the WT, whereas both the *cat2-1 atg1abc* and *cat2-1 atg13ab* mutants phenocopied the *atg1abc* and *atg13ab* mutants, respectively, under carbon starvation conditions (Figure 3G-3I), implying that CAT2 protein itself negatively regulates carbon starvation tolerance by acting upstream of ATG1 and ATG13 in plants.

### CAT2 inhibits ATG1–ATG13 association and autophagy initiation

The ATG1a–ATG13a association is essential for the initiation of autophagy, and we therefore tested whether CAT2 influences this interaction. For the purpose, we first performed yeast-three hybrid (Y3H) assays, where the expression of CAT2 was induced in yeast by the absence of Methionine (Met) when the cells were grown on synthetic dropout (SD) medium. ATG1a interacted with ATG13a in these assays. However, the expression of CAT2 greatly decreased the level of interaction in the yeast cells grown on Met-lacking SD medium (Figure 4A), suggesting that the CAT2 protein interferes with the assembly of the ATG1a-ATG13a complex. We confirmed the role of the CAT2 protein using BiFC assays. cYFP-ATG1a and nYFP-ATG13a were co-expressed with the empty CFP or CFP-CAT2 in *N. benthamiana* leaves. The level of reconstituted YFP fluorescence was significantly weaker in the leaves expressing CFP-CAT2 than in the leaves expressing CFP (Figure 4B, 4C). GST pull-down assays were performed to determine whether CAT2 inhibits ATG1a-ATG13a complex assembly. Both GST-tagged ATG1a and MBP-tagged ATG13a proteins were expressed and purified in *E. coli*, and they were incubated in the absence or presence of the purified 6×His-tagged CAT2 protein *in vitro*. Lower amounts of MBP-ATG13a protein were pulled down by GST-ATG1a in the presence of 6×His-CAT2 than in the absence of 6×His-CAT2 (Figure 4D). This finding further supports the concept that CAT2 has a direct inhibitory action on the ATG1a–ATG13a association.

**Figure 4.**
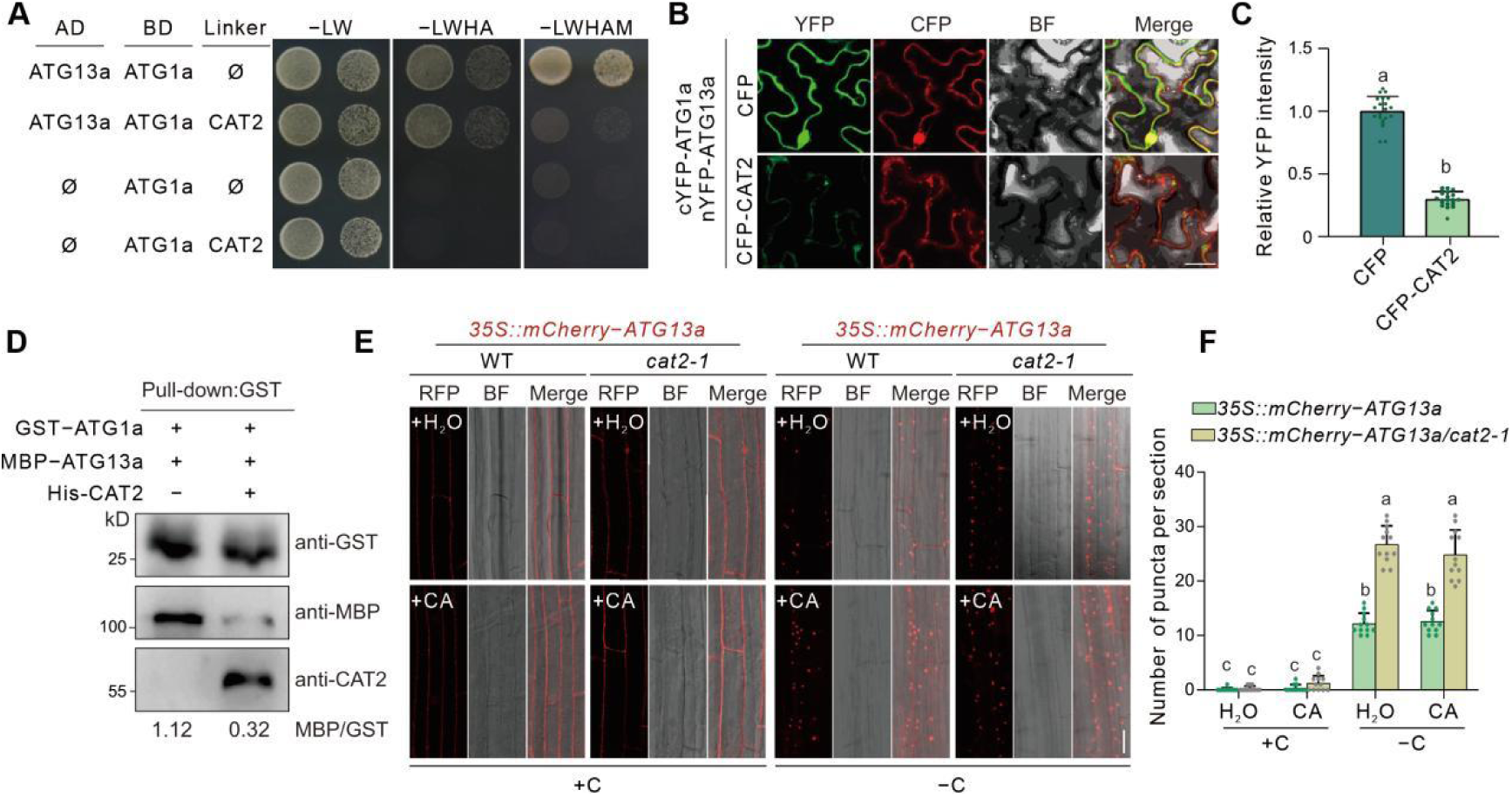
CAT2 inhibits autophagy initiation by dampening ATG1–ATG13 complex formation. (**A**) Yeast-three hybrid assays showing that expression of CAT2 weakens the interaction between ATG1 and ATG13. Yeast cells co-transformed with AD–ATG13a and pBridge–ATG1a or pBridge–ATG1a (CAT2) were dropped onto SD/-LWHA media to validate the interaction of ATG1a with ATG13a. The co-transformed yeast cells were dropped onto SD/-LWHAM medium lacking methionine (Met, M) to induce expression of CAT2. (**B-C**) BiFC assay showing the effect of CAT2 on the interaction between ATG1a and ATG13a. Both cYFP–ATG1a and nYFP–ATG13a were coexpressed with CFP or CFP–CAT2 in *N. benthamiana* leaves for 3 d, and the reconstituted YFP fluorescence was observed (B) and analysed (C) under a confocal microscopy. Bar = 20 μm. Data are mean ± SD (n ≥ 15). Bars with different letters indicate significant differences at p < 0.05, revealed using a one-way analysis of variance with a Tukey’s multiple comparison test. (**D**) *In vitro* GST pull-down assay showing that addition of His–CAT2 reduces MBP–ATG13a binding to GST–ATG1a. Both purified GST–ATG1a and MBP–ATG13a proteins were incubated with the purified His–CAT2, and immobilized on glutathione sepharose beads on ice. After washing, the eluted proteins were subjected to immunoblot analysis with anti-GST, anti-MBP and anti-CAT2 antibodies, respectively. The ratio of MBP–ATG13a to GST–ATG1a was analysed. (**E-F**) Confocal analysis of the *35S::mCherry-ATG13a/WT* and *35S::mCherry-ATG13a/cat2-1* lines. 7-d-old *35S::mCherry-ATG13a/WT* and *35S::mCherry-ATG13a/cat2-1* seedlings grown on sucrose-containing MS medium under long-day conditions were subjected to carbon starvation (–C) conditions in the presence or absence of 1 µM CA for 16 h. The puncta labeled by mCherry-ATG13a was visualized by confocal microscopy (E) and their numbers were analysed (F). Bar = 20 μm. Data are mean ± SD (n ≥ 12). Bars with different letters indicate significant differences at p < 0.05, revealed using a one-way analysis of variance with a Tukey’s multiple comparison test.

Given that the ATG1a-ATG13a association is essential for the initiation of autophagy and subsequent autophagosome formation (Suttangkakul et al., 2011), we investigated whether CAT2 affects autophagy initiation. A 35S::mCherry-ATG13a construct was introduced into WT and *cat2-1* mutant plants, enabling us to observe the autophagy initiation complex with the fluorescence of mCherry-labeled ATG13a. Under carbon sufficient conditions, mCherry fluorescence was evenly distributed in the cytosol in both *35S::mCherry-ATG13a* and *35S::mCherry-ATG13a cat2-1* roots but only weak mCherry fluorescent spots could be detected (Figure 4E, 4F). In contrast, carbon deprivation not only increased the number of mCherry-labeled fluorescent spots, but also the number of spots was greater in the *35S::mCherry-ATG13a cat2-1* roots than in the *35S::mCherry-ATG13a* roots (Figure 4E, 4F). These results indicate that CAT2 inhibits autophagy initiation by disrupting the ATG1a-ATG13a interaction.

### Oxidation of Cys-370 and Cys-413 on the CAT2 protein prevents disruption of ATG1a-ATG13a interactions

While catalases are mainly targeted to peroxisomes, they are synthesized in the cytosol and then transported to the peroxisomes (Zhang et al., 2021, Baker et al., 2023). However, redox-dependent differential accessibility of CAT2 within protein condensates regulates CAT2 compartmentalization between peroxisome, cytosol and nucleus (Lin et al., 2025). To determine the mechanism by which CAT2 regulates autophagy initiation, we generated *35S::GFP-CAT2 cat2-1* plants, in which GFP was fused to the N-terminus of CAT2 to avoid mis-localization (Figure S3A). We observed that GFP-CAT2 was mainly localized in peroxisome-like organelles as well as in cytosol in root cells (Figure S3B). We also generated *35S::GFP-CAT2-SKL cat2-1* transgenic plants, in which a strong type I peroxisome-localization signal peptide, SKL (Ser-Arg-Leu), was fused to the C-terminus of CAT2 (Figure S3A). Indeed, *GFP-CAT2-SKL* was fully localized to peroxisomes, with nearly no detectable cytosolic localization (Figure S3B). In contrast, when WT, *cat2-1*, *35S::GFP-CAT2 cat2-1* and *35S::GFP-CAT2-SKL cat2-1* plants were subjected to carbon starvation, the overexpression of *CAT2* significantly attenuated the starvation tolerance of the *cat2-1* mutants. In contrast, overexpression of *CAT2-SKL* improved the tolerance of the mutants (Figure S3C-F).

We hypothesized that H_2_O_2_ may facilitate autophagy initiation by oxidative modification of the CAT2 protein in the cytosol. We therefore performed BiFC assays using both nYFP-ATG1a and cYFP-ATG13a, together with mCherry-CAT2 or mCherry-SKL as controls in *N. benthamiana* leaves. The level of reconstituted YFP fluorescence in the leaves expressing mCherry-CAT2 was significantly lower than that observed in the leaves expressing mCherry-SKL (Figure 5A-B). Moreover, the addition of H_2_O_2_ significantly enhanced the YFP fluorescence in a concentration-dependent manner in the leaves expressing mCherry-CAT2 but not in the leaves expressing mCherry-SKL. This finding was further confirmed in GST pull-down assays, in which H_2_O_2_ abolished the interactions between CAT2 and ATG1a (Figure 5C) and diminished CAT2-mediated inhibition of the ATG1a-ATG13a interaction (Figure 5D).

**Figure 5.**
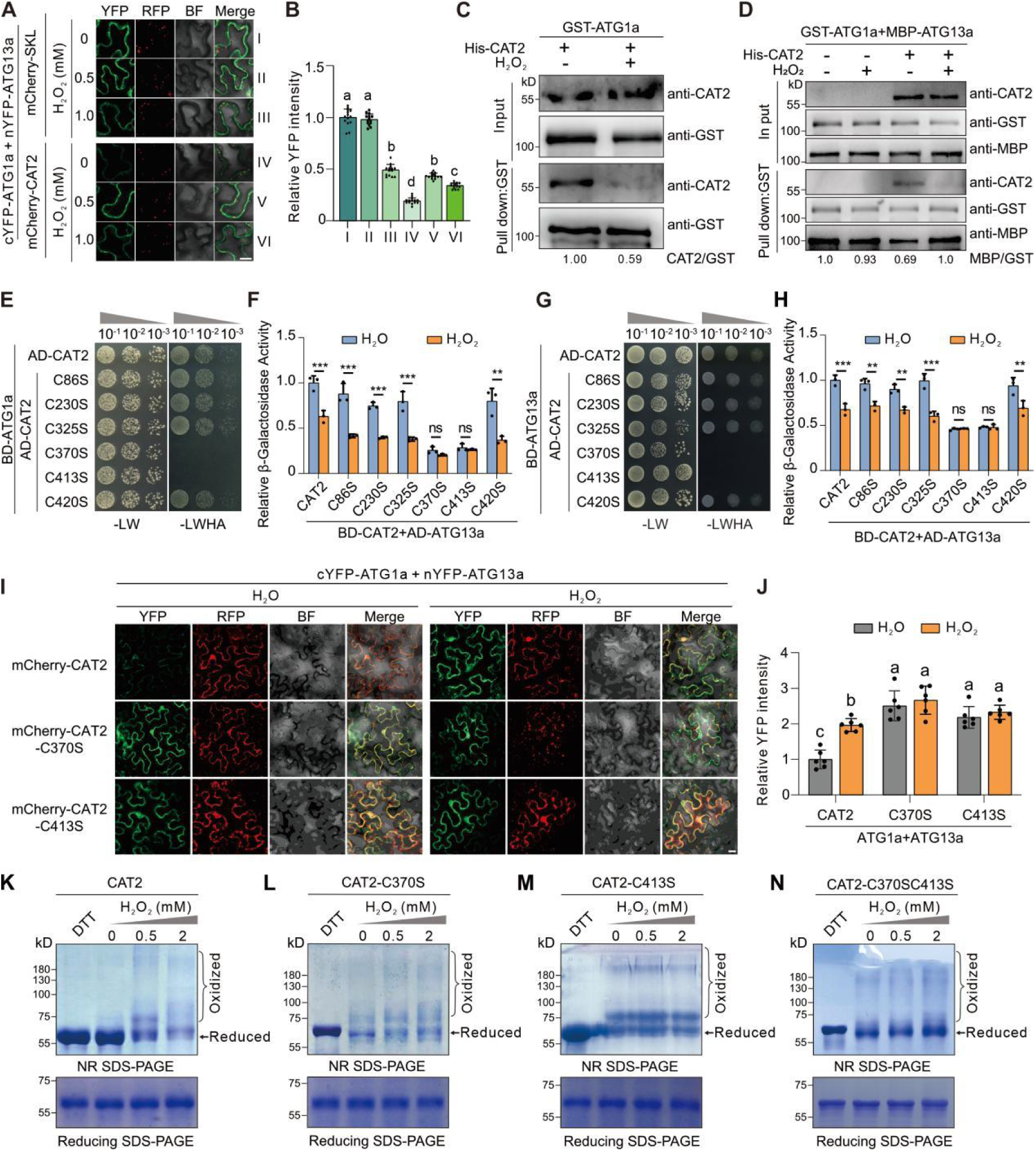
Oxidization of CAT2 at Cys-370 and Cys-413 sites affects its interaction with ATG1a and ATG13a. (**A-B**) BiFC assay showing the role of H_2_O_2_ in CAT2-mediated inhibition of ATG1a-ATG13a interaction. Both cYFP–ATG1a and nYFP–ATG13a were coexpressed with mCherry or mCherry–CAT2 in *N. benthamiana* leaves for 2 d, and then the leaves were treated with 0, 0.5 or 1 mM H_2_O_2_ for 2 h before confocal imaging analysis. The reconstituted YFP fluorescence was observed (A) and analysed (C) under a confocal microscopy. Bar = 20 μm. Data are mean ± SD (n ≥ 15). Bars with different letters indicate significant differences at p < 0.05, revealed using a one-way analysis of variance with a Tukey’s multiple comparison test. (**C**) *In vitro* GST pull-down assay showing the effect of H_2_O_2_ on CAT2 interaction with ATG1a. The purified His–CAT2 protein was first pre-treated with 0 or 0.8 mM H_2_O_2_ at 22°C for 1 h. The mixture of the purified GST–ATG1a and the pre-treated His–CAT2 proteins was immobilized on glutathione sepharose beads at 4°C with rotation for 3 h. After washing with GSH, the eluted proteins were analysed with anti-GST and anti-CAT2 antibodies, respectively. The ratio of His–CAT2 to GST–ATG1a was analysed. (**D**) *In vitro* GST pull-down assay showing the effect of H_2_O_2_ on CAT2-mediated inhibition of ATG1a-ATG13a interaction. The purified His–CAT2 protein was first pre-treated with 0 or 0.8 mM H_2_O_2_ at 22°C for 1 h. The mixture of the purified GST–ATG1a and MBP-ATG13a proteins was incubated with pre-treated His–CAT2 at 4°C with rotation for 3 h, and then immobilized on glutathione sepharose beads. After washing with GSH, the eluted proteins were analysed with anti-GST, anti-MBP, and anti-CAT2 antibodies, respectively. The ratio of MBP-ATG13a to GST–ATG1a was analysed. (**E–H**) Yeast two-hybrid assay assessing the interaction of ATG1a (E) or ATG13a (G) with CAT2 and mutated CAT2, including CAT2-C86S, CAT2-C230S, CAT2-C325S, CAT2-C370S, CAT2-C413S, and CAT2-C420S. The yeast cells cultured in liquid synthetic dropout medium were treated with either H_2_O or 2 mM H_2_O_2_ at 28°C for 1 h. Subsequently, the optical density value of the yeast cell suspension was measured as the normalization reference value, and then the β-galactosidase activity was determined using p-nitrophenyl-β-D-galactoside (ONPG) as the substrate (F, H). Data are mean ± SD (n = 3). Asterisks indicate statistically significant differences (**P<0.01 and ***P<0.001, by ANOVA); “ns” indicates not significant. Bar = 10 mm. (**I–J**) Analysis of the effects of H_2_O_2_ on the interaction between ATG1a and ATG13a in the presence of CAT2, CAT2-C370S or CAT2-C413S using BiFC assays. Both cYFP–ATG1a and nYFP–ATG13a were coexpressed with mCherry–CAT2, mCherry–CAT2-C370S, or mCherry–CAT2-C413S in *N. benthamiana* leaves for 2 d, and then the leaves were treated with either H_2_O_2_ or 0.5 mM H_2_O_2_ for 2 h before confocal imaging analysis. The reconstituted YFP fluorescence was observed under a confocal microscopy (I), and the fluorescence intensities relative to mCherry were analysed (J). Bar = 20 μm. Data are mean ± SD (n = 6). Bars with different letters indicate significant differences at p < 0.05, revealed using a one-way analysis of variance with a Tukey’s multiple comparison test. (**K–N**) Analysis of the purified His-CAT2 proteins treated with DTT and H_2_O_2_. The purified CAT2, CAT2-C370S, CAT2-C413S, and CAT2-C370SC413S were treated with 1 mM DTT or different concentrations of H_2_O_2_ (0, 0.5 or 2 mM) at room temperature for 1 h, and then separated with non-reducing (NR) or reducing SDS-PAGE gels, respectively.

H_2_O_2_ oxidizes the free thiol groups of protein cysteine (Cys, C) residues (Demasi et al., 2021; Liu et al., 2022a). To determine which Cys residues on the CAT2 protein are required for interactions with ATG1a and ATG13a, each of the six Cys in the CAT2 protein, including Cys-86, Cys-230, Cys-370, Cys-413 and Cys-420, were separately mutated to serine (Ser, S) via site-directed mutagenesis (Figure S4A). Y2H experiments were then conducted to determine interactions of the wild-type and mutated forms of the CAT2 protein with ATG1a or ATG13a. CAT2, CAT2-C86S, CAT2-C230S, CAT2-C325S and CAT2-C420S were able to interact with both ATG1a and ATG13a in yeast cells, whereas no interactions were observed with CAT2-C370S and CAT2-C413S (Figure 5E, 5G). Next, we treated the yeast cells with either H_2_O or H_2_O_2_ and measured their β-glactosidase activity to quantify the effects of H_2_O_2_ on the CAT2 interaction with ATG1a and ATG13a. The presence of H_2_O_2_ significantly weakened the interactions between ATG1a or ATG13a and CAT2, CAT2-C86S, CAT2-C230S, CAT2-C325S and CAT2-C420S (Figure 5F, 5H). However, this effect was not observed with CAT2-C370S and CAT2-C413S (Figure 5F, 5H). The finding that H_2_O_2_ prevents the CAT2 interaction with ATG1a and ATG13a by oxidant Cys-370 and Cys-413, was supported by other studies showing that expression of mCherry-CAT2 but not mCherry-CAT2-C370S or mCherry-CAT2-C413S markedly reduced the interactions between ATG1a and ATG13a in *N. benthamiana* leaves. In addition, the mCherry-CAT2-mediated inhibition of ATG1a-ATG13a interaction was disrupted in the presence of H_2_O_2_ but there was no effect with mCherry-CAT2-C370S or mCherry-CAT2-C413S (Figure 5I, 5J). These findings support the conclusion that H_2_O_2_ -mediated oxidation of Cys-370 and Cys-413 on the CAT2 protein prevents the inhibition of ATG1a-ATG13a interactions.

### Oxidation of CAT2 Cys-370 and Cys-413 improves carbon starvation tolerance

We purified 6×His-tagged CAT2, CAT2-C370S, CAT2-C413S and CAT2-C370SC413S recombinant proteins in *E. coli*, to investigate whether Cys-370 and Cys-413 in CAT2 are directly oxidized by H_2_O_2_. We then added the reducing agent dithiothreitol (DTT) or H_2_O_2_ *in vitro* and separated the proteins on nonreducing and reducing SDS-PAGE gels, respectively. Under denaturing non-reducing conditions, both DTT-treated and untreated CAT2 proteins migrated predominantly as a monomeric band. Following H_2_O_2_ treatment, however, multiple oxidized protein bands appeared on the gels, while the reduced protein bands were decreased (Figure 5K). Although untreated CAT2-C370S, CAT2-C413S and CAT2-C370SC413S proteins exhibited a multi-band profile resembling oxidized CAT2, the addition of H_2_O_2_ exposure did not alter the pattern of migration on the gels (Figure 5L-N), implying that both Cys-370 and Cys-413 are the major sites for H_2_O_2_-mediated oxidization of the CAT2 protein.

To study the role of Cys-370 and Cys-413 in CAT2-mediated regulation of autophagy in plants, we complemented the *cat2-1* mutant by expressing *CAT2*, *CAT2-C370S* and *CAT2-C413S* under the control of the native *CAT2* promoter, resulting in *CAT2::CAT2 cat2-1*, *CAT2::CAT2-C370S cat2-1*, and *CAT2::CAT2-C413S cat2-1* plants, respectively. The expression of *CAT2*, *CAT2-C370S* and *CAT2-C413S* restored the absence of CAT2 protein and the low catalase activity in the *cat2-1* mutants to levels similar to those in the WT (Figure S4B, S4C), indicating that C-S substitutions of Cys-370 or Cys-413 retain CAT2 catalase activity. We next measured the autophagosomes in the WT, *cat2-1*, *CAT2::CAT2 cat2-1*, *CAT2::CAT2-C370S cat2-1*, and *CAT2::CAT2-C413S cat2-1* plants. Under carbon deprivation conditions, the *cat2-1* mutant exhibited more autophagosomes than the WT, but the *CAT2::CAT2 cat2-1* plants had a similar number of autophagosomes to the WT (Figure 6A, 6B). In contrast, the number of autophagosomes was significantly higher in the *CAT2::CAT2-C370S cat2-1* and *CAT2::CAT2-C413S cat2-1* plants than in the WT or the *CAT2::CAT2 cat2-1* plants (Figure 6A, 6B), This finding suggests that both Cys-370 and Cys-413 are required for CAT2-mediated inhibition of autophagy in Arabidopsis. Consistent with this finding, the expression of *CAT2* fully complemented the carbon starvation phenotypes of the *cat2-1* mutant in both the early and the late stages of carbon deprivation. In contrast, the *CAT2::CAT2-C370S cat2-1* and *CAT2::CAT2-C413S cat2-1* plants showed a higher tolerance to carbon starvation than the WT and *CAT2::CAT2 cat2-1* plants at both the early and the late stages (Figure 6C-6E). These findings confirm that Cys-370 and Cys-413 are required for the CAT2-mediated inhibition of carbon starvation tolerance in Arabidopsis. Taken together, these results demonstrate that H_2_O_2_-mediated oxidation of Cys-370 and Cys-413 on the CAT2 protein regulates the inhibition of autophagy and so improves carbon starvation tolerance (Figure 6F).

**Figure 6.**
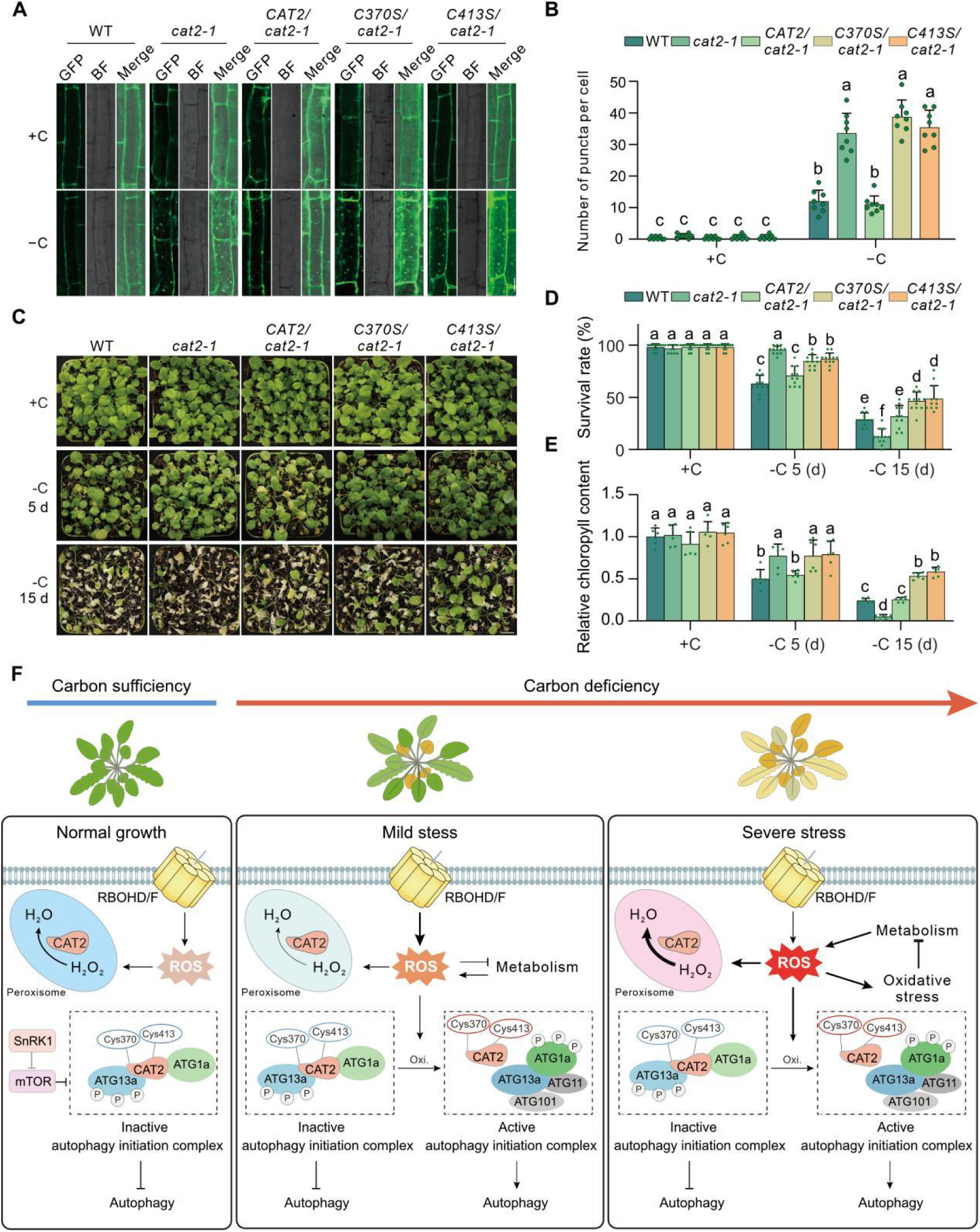
Analysis of Autophagy Levels in CAT2/cat2, CAT2-C370S/cat2 and CAT2-C413S/cat2. (**A–B**) Autophagosomes in the WT, *cat2-1, CAT2/cat2-1, C370S/cat2-1,* and *C413S/cat2-1* lines using MDC staining. 7-d-old WT, *cat2-1, CAT2::CAT2 cat2-1* (*CAT2/cat2-1*)*, CAT2::CAT2-C370S cat2-1* (*C370S/cat2-1*), and *CAT2::CAT2-C413S cat2-1* (*C413S/cat2-1*) lines were subjected to carbon starvation (–C) condition in the absence or presence of 1 µM CA for 16 h, and the autophogosomes stained with MDC were observed under a confocal microscopy (A), and the number of puncta per cell was analysed (B). Bar = 20 μm. Data are mean ± SD (n ≥ 7). Bars with different letters indicate significant differences at p < 0.05, revealed using a one-way analysis of variance with a Tukey’s multiple comparison test. (**C–E**) Phenotypes (C), survival rates (D) and relative chlorophyll content (E) of soil-grown WT, *cat2-1, CAT2/cat2-1, C370S/cat2-1,* and *C413S/cat2-1* plants treated with carbon starvation for 0, 5 and 15 d. Data are means ± SD (n ≥ 6). Bars with different letters indicate significant differences at P < 0.05, revealed using a one-way analysis of variance with a Tukey’s multiple comparison test. (**F**) A working model for CAT2/H_2_O_2_-mediated modulation of autophagy during carbon starvation in Arabidopsis. Schematic model summarizes the proposed dynamics from carbon-sufficiency/normal growth (left) to carbon-deficiency, including the early stage/mild stress (middle) and the late stage/severe stress (right) conditions. Under normal conditions, H_2_O_2_ levels are low, while cytosolic CAT2, in a reduced state, interacts with both ATG1 and ATG13 to inhibit the assembly of autophagy initiation complex. During the early stage of carbon starvation, increased H_2_O_2_ oxidizes cytosolic CAT2 at its Cys-370 and Cys-413 residues, dampening its interaction with both ATG1 and ATG13, facilitating autophagy initiation complex assembly and the activation of autophagy. In the late stage of carbon starvation, overaccumulated H_2_O_2_ elicits oxidative injury and erodes plant viability, thus peroxisomal CAT2 is necessary to counteract cellular oxidative stress by eliminating excess H_2_O_2_, even though higher H_2_O_2_ may still have a role in regulating autophagy via oxidizing CAT2 in the cytosol.

## Discussion

Autophagy has fundamental roles in plant adaptive responses to environmental stresses, such as carbon starvation, and is precisely regulated by the TOR-SnRK1-mediated energy sensing pathway (Suttangkakul et al., 2011). This pathway is also regulated by ROS, which regulate plant growth, development and stress responses (Considine and Foyer, 2023, 2024; Liu et al., 2025). However, the precise roles of ROS in the regulation of autophagy have remained largely unresolved (Zhang et al., 2026). In this study, we demonstrate the presence of a new mechanism by which carbon starvation-mediated H_2_O_2_ accumulation regulates autophagy and carbon starvation tolerance in plants. We present data showing that the CAT2 protein interacts with both ATG1 and ATG13 in the cytosol in the absence of stress, and that this interaction prevents ATG1-ATG13 complex assembly, thus inhibiting autophagy initiation. The accumulation of H_2_O_2_ during the early stage of carbon starvation releases the CAT2-dependent inhibition of autophagy initiation. This regulation is achieved through the H_2_O_2_-dependent oxidation of the Cys-370 and Cys-413 residues on the CAT2 protein in the cytosol, preventing the inhibition of the ATG1 and ATG13 association. At the later stages of carbon starvation, CAT functions to limit chlorosis (Figure 6F).

### CAT2 is an autophagy-regulation factor

This study has revealed that CAT2 acts as a redox-regulated molecular switch that couples H_2_O_2_ signaling to autophagosome formation. In the absence of starvation stress, the formation of the ATG1-ATG13 initiation complex is prevented by association with CAT2, which structurally inhibits ATG1-ATG13 interactions. In the cytosol CAT2 is likely to be present largely in a monomer form that lacks activity (Li et al., 2015). The maturation of active CAT enzyme involves the insertion of a heme group (Fe-protoporphyrin IX) into the apo-monomer but there is much uncertainty about where the tetrameric CAT protein is formed (Baker et al., 2023).

The data presented here shows that the interaction of CAT2 with ATG1a or ATG13a occurs in the cytosol and not in the peroxisomes or autophagosomes (Figure 3B and 5A). The importance of CAT as an antioxidant enzyme is well established. Furthermore, accumulating evidence has demonstrated that CAT is a moonlighting protein that can serve very different functions, in addition to its catalytic role (Baker et al., 2023). We have previously shown that the relocation of CAT to the nucleus occurs in a ROS-dependent manner (Lin et al., 2025). Moreover, CAT isoforms are targets of many pathogen effectors, in a manner that regulates programmed cell death. Different pathogen effectors manipulate CAT transcription (Zhu et al., 2023), activity (Mathioudakis et al., 2013; Roshan et al., 2018), stability (Zhang et al., 2015; You et al., 2022) and location (Zhang et al., 2015).

We present data showing that the purified CAT2 protein can be oxidized by the addition of exogenous H_2_O_2_, forming both monomers and oligomers (Figure 5K). Furthermore, we have demonstrated that Cys-370 and Cys-413 are oxidized in the presence of H_2_O_2_, and that substitution of C370S and C413S with other amino acids blocks the oxidation of the CAT2 protein (Figure 5L-5N). In addition, neither CAT2-C370S nor CAT2-C413S could interact with ATG1 and ATG13, and CAT2 but not CAT2-C370S or CAT2-C413S restored autophagosome numbers in the *cat2-1* mutants to WT levels under carbon starvation conditions (Figure 6A). These findings support the conclusion that Cys-370 and Cys-413 are key sites for the regulation of CAT2 interactions with ATG1 and ATG13.

### The role of H_2_O_2_ in carbon starvation

The data presented here suggest that H_2_O_2_ fulfils multiple functions in the carbon starvation response. The activation of RBOH enzyme activity at the initial phase of the starvation response (Figure S1A-D) promotes carbon starvation tolerance by initiating autophagy, through the formation of the ATG1-ATG13 autophagy initiation complex. This regulation was not observed in the *atg1abc* or *atg13ab* mutants (Figure 2A-B). The *rbohd/f* mutant with lower H_2_O_2_ levels exhibited more pronounced chlorosis at the early stages of carbon starvation (e.g. 3 days) but had poor levels of survival at the later stages of carbon starvation (e.g. 8 days) compared to the WT (Figure 1). The addition of H_2_O_2_ restored the carbon starvation sensitive phenotype in the *rbohd/f* mutants (Figure 1). Conversely, the *cat2-1* mutants had higher levels of autophagy initiation complex, autophagic flux and less pronounced chlorosis at the early stages of carbon starvation (e.g. 3 days) (Figure 1 and 2F-I). The level of DAB staining was higher at the later stages of carbon starvation than the early stages (Figure 1A-B). It is likely, that high ROS levels at the later stages of starvation trigger other genetic programs such as programmed cell death, depending on what redox sensing proteins are produced at that time point. The *cat2-1* mutant exhibited more chlorosis with poorer survival rates than the WT (Figure 1 and 2F). The removal of H_2_O_2_ through the application of the reducing agent potassium iodide (KI) significantly increased the carbon starvation tolerance of the *cat2-1* mutant (Figure 1). These findings suggest that H_2_O_2_ fulfils other signaling roles in the regulation of in plant autophagy that operate in a time-dependent manner.

The H_2_O_2_-dependent oxidization of CAT2 at the Cys-370 and Cys-413 sites, releases inhibition of ATG1-ATG13 complex, facilitating autophagy in plants. The Cys residue at position 370 (Cys-370) is highly conserved, whereas the Cys residue at position 413 (Cys-413) exhibits variability in different plant species (Figure S4D, E). Together, these findings suggest that H_2_O_2_-mediated oxidation of CAT2 at Cys-370 is a conserved mechanism for the regulation of autophagy initiation and carbon starvation responses in plants.

### Divergent and convergent signaling pathways and mechanisms underlying autophagy

In animals, H_2_O_2_ directly oxidizes the cysteine residue at position 81 (Cys-81) in the core autophagy protein HsATG4A, thereby inhibiting its proteolytic activity and regulating the ATG8-PE binding process (Scherz-Shouval et al., 2007). This oxidation-dependent regulatory mechanism, which relies on the conserved Cys residue in ATG4, has been further validated in yeast and Chlamydomonas (Perez-Perez et al., 2016; Perez-Perez et al., 2014). Sequence alignment in plants indicates that the homologous site corresponding to human ATG4’s Cys-81 is a serine residue instead of a cysteine (Zhang et al., 2026), excluding the possibility that H_2_O_2_ modulates autophagy by oxidizing ATG4 at this site in plants. This critical difference clearly suggests that plants are likely to employ a unique strategy distinct from animals to transmit ROS signals to the autophagy pathway.

This study establishes the H_2_O_2_/CAT2-ATG1a-ATG13a module as a key regulatory node in the initiation of cellular autophagy. We found that ROS can directly regulate the ATG1-ATG13 initiation complex by acting on CAT2, thereby manipulating the initiation of autophagy under carbon starvation stress. This finding, together with previously reported regulatory mechanisms, such as GRF6/8-dependent SINAT-mediated ubiquitination and degradation of ATG13 under carbon starvation stress, TOPP-mediated dephosphorylation of ATG13 under carbon starvation stress, and isolation of key autophagy proteins including ATG1 and ATG13 by heat shock bodies under heat stress (Li et al., 2024; Qi et al., 2022; Wang et al., 2022), collectively demonstrates that the ATG1-ATG13 autophagy initiation complex serves as a central hub converging multiple upstream signals to precisely regulate autophagy initiation.

This study demonstrates the presence of a H_2_O_2_-CAT2 switch that regulates autophagy. We demonstrate that CAT2 directly inhibits ATG1a-ATG13a complex formation in the cytosol in a redox-dependent manner in the absence of stress. The H_2_O_2_-oxidation of CAT2 is initiated by carbon starvation and this facilitates the ATG1a-ATG13a interaction and initiates autophagy. Other ROS-regulated processes occur at the later stages of carbon starvation. Further research is required to determine what redox-regulated processes are involved in the regulation of chlorosis and programmed cell death.

## MATERIALS AND METHODS

### Plant materials

Arabidopsis (*A. thaliana*) ecotype Columbia-0 was used as the WT. The *cat2-1* (SALK_076998), *rbohd* (SALK_120299) and *rbohf* (SALK_034674) mutants were obtained from the Arabidopsis Biological Resource Center. The seeds of *atg1abc*, *atg13ab* and *GFP-ATG8e* were previously reported (Qi et al., 2022), and verified by genomic DNA PCR. The *cat2-1 atg1abc* quadruple mutant and *cat2-1 atg13ab* triple mutant were obtained by genetic crossing. The transgenic plants *CAT2pro::CAT2 cat2-1*, *CAT2pro::CAT2-C370S cat2-1*, *CAT2pro::CAT2-C413S cat2-1*, *35S::GFP-CAT2 cat2-1*, *35S::GFP-CAT2-SKL cat2-1* and *mCherry-ATG13a* were generated as described below in methods. Subsequently, the *mCherry-ATG13a cat2-1*, *GFP-ATG8e atg13ab* and *GFP-ATG8e atg13ab cat2-1* plants were obtained by genetic crossing.

Seeds were surface sterilized using 10% (w/v) sodium hypochlorite (NaClO) for 5 min, washed 5 times with sterile water, and sown on a 1/2× MS medium (pH 5.8, containing 1% [w/v] sucrose and 0.8% [w/v] agar). Plates were kept at 4 °C for 3 d, and then seeds were germinated and grown under a 16 h/8 h light/dark photoperiod at 100 μmol m^−2^ s^−1^ at 22 °C.

For carbon starvation treatment on MS medium, 1-week-old seedlings grown on 1/2× MS were transferred to solid sucrose-free medium and incubated under continuous dark conditions until the seedlings started showing poor growth. After recovery under normal growth conditions for 7 days, seedling phenotypes were recorded as photographs, and the survival rate was calculated as the percentage of seedlings with obvious regreening and the appearance of new leaves.

For carbon starvation treatment in soil, 1-week-old seedlings grown on 1/2× MS were transplanted in soil for additional 2 weeks under a 16 h/8 h light/dark photoperiod at 100 μmol m^−2^ s^−1^ at 22 °C, and then the plants were incubated under continuous dark conditions for about 7-10 d. After recovery under normal growth conditions for 7 days, seedling phenotypes were recorded as photographs, and the survival rate was calculated as the percentage of plants with obvious regreening and the appearance of new leaves.

For biochemical treatments, 1-week-old Arabidopsis seedlings grown on 1/2× MS medium containing 1% (w/v) sucrose were transferred to either solid MS medium or sucrose-free solid MS medium (–C) containing 1 μM concanamycin A (CA) for treatment, followed by total protein extraction and immunoblot analysis.

### Plasmid construction and plant transformation

The pCambia1300 vector was digested at *Nco*I and *Pst*I sites to obtain a modified vector without the 35S promoter and GFP. To generate the 1300-CAT2pro:CAT2 transgenic lines, the genomic sequence containing 1997 bp upstream of the *CAT2* translation start codon (ATG) was amplified and cloned into the modified pCambia1300 vector at the *Pst*1 site, resulting in 1300-CAT2pro plasmid. Then, the full length CDSs of *CAT2*, *CAT2-C370S* and *CAT2-C413S* were amplified using PCR and cloned into the 1300-CAT2pro plasmid behind the *CAT2* promoter, resulting in 1300-CAT2pro::CAT2, 1300-CAT2pro::CAT2-C370S and 1300-CAT2pro::CAT2-C413S plasmids, respectively. The full length CDS of *ATG13a* were amplified using PCR and cloned into pmCherry vector, generating 35S::ATG13a-mCherry plasmid. The primers used for plasmids construction were listed in Supplemental Table S1.

The resultant plasmids were introduced into the WT or *cat2-1* mutant plants via an Agrobacterium (GV3101)-mediated transformation and the floral dip method. Transformants were selected on antibiotic medium and single-copy transgenic lines were selected. T4 homozygous transgenic plants were used in this study.

### Y2H assays

pGADT7-CAT2 constructed was previously reported (Zhang et al., 2021). The full-length CDSs of *ATG1a*, *ATG1b*, *ATG1c*, *ATG13a* and *ATG13b* were cloned into pGBKT7 (Clontech) at the *BamH*I site, respectively. Yeast transformation and growth were carried out using the Matchmaker system (Clontech) according to the manufacturer’s protocols. Yeast transformants were selected on double dropout medium lacking Leu and Trp (−LW), and protein interactions were analyzed on quadruple dropout medium lacking Leu, Trp, His, and Ade (−LWHA). Primer sequences are listed in Supplemental Table S1.

### BiFC assay

The CDSs of *CAT2*, *ATG1a*, and *ATG13a* were cloned into the pSPYCE or pSPYNE vector (Walter et al., 2004) containing the C-terminal half of YFP (cYFP) and or the N-terminal half of YFP (nYFP), respectively. The peroxisomal marker mCherry-SKL (Yuan et al., 2017), nYFP-CAT2 and cYFP-ATG13a or cYFP-ATG1a were co-expressed in *N. benthamiana* leaves for 3 d, after which YFP (excitation 488 nm and emission 520 to 560 nm) and mCherry (excitation 561 nm and emission 600 to 630 nm) fluorescence signals were detected using a laser scanning confocal microscope (Zeiss LSM980, Zeiss, Germany). Primer sequences are listed in Supplemental Table S1.

### Co-IP assays

To perform Co-IP assays, the CDS of *ATG1a* was cloned into pCambia1300 at the *BamH*I site, including sequences encoding a GFP tag fused to its C terminus and under the control of the *35S* promoter. The Co-IP experiment was performed according to the previously reported methods with some modifications (Lu et al., 2023; Zhang et al., 2025). Briefly, GFP-tagged ATG1a and MYC-tagged CAT2 were transformed into Agrobacterium GV3101 and infiltrated into 3-week-old *N. benthamiana* leaves. After 3 d, total proteins were extracted from the infected leaves by homogenization in IP buffer (150 mM NaCl, 25 mM Tris-HCl, pH 7.5, 0.2% Nonidet P-40, 1 mM phenylmethylsulfonyl fluoride, and 1× protease inhibitor cocktail) and then immunoprecipitated by an anti-GFP antibody (ABclonal, AE012, diluted 1:1,000). The resulted precipitates were resuspended and detected using an anti-GFP (ABclonal, AE012) and anti-CAT2 (Zhang et al., 2020) antibodies, respectively.

### GST pull-down assay

The CDSs of full-length *CAT2* and *ATG1a* were cloned into the pET28a (6×His) and pGEX4T-1 vectors, respectively. The CDSs of full-length *CAT2* and *ATG13a* were cloned into pEGX4T-1 vectors and pAB-6×His-MBP vectors, respectively, and subjected to GST pull-down experiments. The GST pull-down assay was performed according to our previous reports (Liu et al., 2022b; Lu et al., 2023). Briefly, GST-ATG1a and 6×His-CAT2 were expressed and purified in *E. coli* strain BL21 (DE3). 6×His-CAT2 was mixed with GST-ATG1a or GST alone on ice for 1 h and then incubated with GST-Sefinose Resin 4FF (Settled Resin) (Sangon Biotech, C600031) at 4 °C for 3 h. After washing, the eluted protein eluted with Elution Buffer (10 mM GSH in 50 mM Tris-HCl, pH 8.0) was detected with anti-His (Beyotime, AH306) and anti-GST (ABclonal, AE006) antibodies, respectively. The purified GST-CAT2 and MBP-ATG13a proteins were incubated as shown above, and the final eluted proteins were detected with anti-MBP (Beyotime, #AF2912) and anti-GST (ABclonal, AE006) antibodies, respectively. Primer sequences are listed in Supplemental Table S1.

### Yeast three-hybrid assays

Yeast three-hybrid assays were performed following previous descriptions (Tan et al., 2023). Briefly, bait construct BD-ATG13a, prey construct AD-ATG1a, and pBridge-ATG1a (CAT2) were co-transformed into Y190 yeast cells while transformants were screened on the selective medium SD/−Leu/−Trp/−His (−LWH). Positive clones were picked and resuspended in sterile water. The suspensions were serially diluted. Each dilution was then spotted onto SD/−Leu/−Trp (−LW), SD/−Leu/−Trp/−His (−LWH), and SD/−Leu/−Trp/−His/-Met (−LWHM) plates for growth assessment. β-Galactosidase activities were measured as described previously by liquid culture assays using 2-Nitrophenyl β-D-galactopyranoside (ONPG) as the substrate according to Yeast Protocols Handbook (Clontech) (Tan et al., 2023).

### Detection of catalase activity

The seedlings treated with or without carbon starvation were ground to a fine powder under liquid nitrogen and suspended in freshly prepared cold protein extraction buffer (50 mM potassium phosphate buffer, pH 7.8, 0.2 mM ethylenediaminetetraacetic acid (EDTA)-Na_2_, 0.1 mM ascorbic acid, and 1% polyvinyl polypyrrolidone, PVPP). After centrifugation at 12,000 *g* for 10 min at 4°C, the supernatant was transferred and used to detect the catalase activity. The catalase activity was determined according to the published methods by monitoring the consumption of H_2_O_2_ at 240 nm (Aebi, 1984).

### Detection of RBOH activity

RBOH activity was determined according to the previously reported method (Jasso-Robles et al., 2020). Briefly, protein extracts (300 μg) were resolved electrophoretically in a native 10 % polyacrylamide gel at 120 V. Then the gel was rinsed in 10 mM Tris-HCl (pH 7.4) for 30 min. Afterwards, the gel was incubated in the dark for 30 min in a reaction mixture containing 10 mM Tris-HCl buffer (pH 7.4), 1 mM EDTA, 0.5 mg/ml NBT, and 134 μM NADPH, until the color bands were clearly visible. Band intensities were quantified with Image J program. RBOH activities were expressed as relative units (RU).

### Detection of CAT2 protein oligomerization and oxidation

To assay the oxidation and oligomerization of CAT2, purified CAT2, CAT2-C370S, CAT2-C413S and CAT2-C370SC413S proteins were treated with different concentrations of H_2_O_2_ or 1 mM DTT for 60 min at room temperature, then separated on reducing or non-reducing SDS-PAGE gels, respectively. The proteins were visualized using Coomassie Brilliant Blue staining.

### DAB staining

7-d-old WT, *cat2-1* and *rbohd/f* seedlings were treated with carbon starvation for 0 ∼ 7 d under dark, then their H_2_O_2_ accumulation was assayed using DAB staining, as described previously (Liu et al., 2022a; Zhang et al., 2021). Briefly, the seedlings were incubated in freshly prepared DAB staining solution (1 mg/ml DAB and 0.1% Tween 20 in 10 mM Na_2_HPO_4_) for 6 h, and then rinsed with 70% ethanol to remove the chlorophyll.

### Statistical analysis

Data are means ± SD of 3 biological replicates, and the asterisks indicate statistically significant differences (*P < 0.05, **P < 0.01, and ***P < 0.001, Student’s *t* test). Bars with different letters indicate significant differences at P < 0.05 by ANOVA with Tukey’s multiple comparison test.

### Accession numbers

Sequence data from this article can be found in the Arabidopsis Genome Initiative or GenBank/EMBL databases under the following accession numbers: *CAT2* (AT4G35090), *ATG1a* (AT3G61960), *ATG1b* (AT3G53930), *ATG1c* (AT2G37840), *ATG13a* (AT3G49590), *ATG13b* (AT3G18770), *ATG8e* (AT2G45170) and *ACTIN2* (AT3G18780).

## ACKNOWLEDGMENTS

We are grateful to Prof. Shi Xiao (Sun Yat-sen University, China) and Prof. Xingliang Hou (South China Botanical Garden, Chinese Academy of Sciences, China) for providing the *atg13ab*, *atg1abc*, *GFP-ATG8e* seeds. This work was supported by the National Natural Science Foundation of China (#32570340 and #32322010), Scientific Research Innovation Capability Support Project for Young Faculty (SRICSPYF-BS2025099), and the Natural Science Foundation of Henan Province (252300421075). CHF thanks the Leverhulme Trust (RPG2021-126) for financial support.

## AUTHOR CONTRIBUTIONS

W.-C.L and C.-H.F conceived the work. C.Z., S.-Q.L., P.J., J.-X.W., B.-H-C., R.-F.S., C.-Y.L K.-K-L., and H.Y conducted the experiment. C.Z. analyzed the data and prepared the figures. W.-C.L and C.-H.F wrote the paper. All authors read the manuscript.

## REFERENCES

Aebi, H. 1984. Catalase in vitro. Methods Enzymol 105, 121–126.

Al-Hajaya, Y. A.-R., Karpinska, B., Foyer, C.H., Baker, A. 2021. Nuclear and peroxisomal localisation and functional characterisation of Arabidopsis CATALASE2 C terminal variants. Plant Cell Environ 45, 1096–1108.

Baker. A., Lin, C.-C., Lett, C., Karpinska, B., Wright, M.H., Foyer, C.H. 2023. Catalase: a critical node in the regulation of cell fate. Free Rad. Biol. Med. 199, 56–66.

Considine, M. J., Foyer, C.H. 2023. Metabolic regulation of quiescence in plants. Plant J. 114, 1132–1148.

Considine, M.J., Foyer, C.H. 2024. Redox regulation of meristem quiescence: Outside/ in. J. Expt Bot, 75, 6037–6046

Demasi, M., Augusto, O., Bechara, E.J.H., Bicev, R.N., Cerqueira, F.M., da Cunha, F.M., Denicola, A., Gomes, F., Miyamoto, S., Netto, L.E.S., Randall, L.M., Stevani, C.V., Thomson, L. 2021. Oxidative Modification of Proteins: From Damage to Catalysis, Signaling, and Beyond. Antioxid Redox Signal 35(12), 1016–1080.

Fu, Z.W., Feng, Y.R., Gao, X., Ding, F., Li, J.H., Yuan, T.T., Lu, Y.T. 2023. Salt stress-induced chloroplastic hydrogen peroxide stimulates pdTPI sulfenylation and methylglyoxal accumulation. Plant Cell 35(5), 1593–1616.

Jasso-Robles, F.I., Gonzalez, M.E., Pieckenstain, F.L., Ramirez-Garcia, J.M., Guerrero-Gonzalez, M.L., Jimenez-Bremont, J.F., Rodriguez-Kessler, M. 2020. Decrease of Arabidopsis PAO activity entails increased RBOH activity, ROS content and altered responses to Pseudomonas. Plant Sci 292, 110372.

Kurusu, T., Koyano, T., Hanamata, S., Kubo, T., Noguchi, Y., Yagi, C., Nagata, N., Yamamoto, T., Ohnishi, T., Okazaki, Y., Kitahata, N., Ando, D., Ishikawa, M., Wada, S., Miyao, A., Hirochika, H., Shimada, H., Makino, A., Saito, K., Ishida, H., Kinoshita, T., Kurata, N., Kuchitsu, K. 2014. OsATG7 is required for autophagy-dependent lipid metabolism in rice postmeiotic anther development. Autophagy 10(5), 878–888.

Li, X., Liao, J., Chung, K.K., Feng, L., Liao, Y., Yang, Z., Liu, C., Zhou, J., Shen, W., Li, H., Yang, C., Zhuang, X., Gao, C. 2024. Stress granules sequester autophagy proteins to facilitate plant recovery from heat stress. Nat Commun 15(1), 10910.

Lin, C.-C., Foyer, C.H., Wright, M., Baker, A. 2025. Redox regulation of LSD1/CATALASE 2 phase separation condensates controls location and functions. New Phytol. 247, 2824–2838.

Liu, W.C., Halliwell, B., Foyer, C.H. 2025. The critical importance of accurate chemical notation for the superoxide radical (O (2) (*-)) in the plant literature. Nat Plants.

Liu, W.C., Han, T.T., Yuan, H.M., Yu, Z.D., Zhang, L.Y., Zhang, B.L., Zhai, S., Zheng, S.Q., Lu, Y.T. 2017. CATALASE2 functions for seedling postgerminative growth by scavenging H(2) O(2) and stimulating ACX2/3 activity in Arabidopsis. Plant Cell Environ 40(11), 2720–2728.

Liu, W.C., Song, R.F., Qiu, Y.M., Zheng, S.Q., Li, T.T., Wu, Y., Song, C.P., Lu, Y.T., Yuan, H.M. 2022a. Sulfenylation of ENOLASE2 facilitates H(2)O(2)-conferred freezing tolerance in Arabidopsis. Dev Cell 57(15), 1883–1898 e1885.

Liu, W.C., Song, R.F., Zheng, S.Q., Li, T.T., Zhang, B.L., Gao, X., Lu, Y.T. 2022b. Coordination of plant growth and abiotic stress responses by tryptophan synthase beta subunit 1 through modulation of tryptophan and ABA homeostasis in Arabidopsis. Mol Plant 15(6), 973–990.

Lu, K.K., Song, R.F., Guo, J.X., Zhang, Y., Zuo, J.X., Chen, H.H., Liao, C.Y., Hu, X.Y., Ren, F., Lu, Y.T., Liu, W.C. 2023. CycC1;1–WRKY75 complex-mediated transcriptional regulation of SOS1 controls salt stress tolerance in Arabidopsis. The Plant Cell 35(7), 2570–2591.

Michelet, L., Roach, T., Fischer, B.B., Bedhomme, M., Lemaire, S.D., Krieger-Liszkay, A. 2013. Down-regulation of catalase activity allows transient accumulation of a hydrogen peroxide signal in Chlamydomonas reinhardtii. Plant Cell Environ 36(6), 1204–1213.

Minina, E.A., Moschou, P.N., Vetukuri, R.R., Sanchez-Vera, V., Cardoso, C., Liu, Q., Elander, P.H., Dalman, K., Beganovic, M., Lindberg Yilmaz, J., Marmon, S., Shabala, L., Suarez, M.F., Ljung, K., Novak, O., Shabala, S., Stymne, S., Hofius, D., Bozhkov, P.V. 2018. Transcriptional stimulation of rate-limiting components of the autophagic pathway improves plant fitness. J Exp Bot 69(6), 1415–1432.

Mittler, R., 2017. ROS Are Good. Trends in Plant Science 22(1), 11–19.

Mittler, R., Vanderauwera, S., Suzuki, N., Miller, G., Tognetti, V.B., Vandepoele, K., Gollery, M., Shulaev, V., Van Breusegem, F. 2011. ROS signaling: the new wave? Trends Plant Sci 16(6), 300–309.

Nukarinen, E., Nägele, T., Pedrotti, L., Wurzinger, B., Mair, A., Landgraf, R., Börnke, F., Hanson, J., Teige, M., Baena-Gonzalez, E., Dröge-Laser, W., Weckwerth, W., 2016. Quantitative phosphoproteomics reveals the role of the AMPK plant ortholog SnRK1 as a metabolic master regulator under energy deprivation. Scientific Reports 6(1), 31697.

Otegui, M., Soto-Burgos, J., Bassham, D.C. 2017. SnRK1 activates autophagy via the TOR signaling pathway in Arabidopsis thaliana. Plos One 12(8), e0182591.

Perez-Perez, M.E., Lemaire, S.D., Crespo, J.L. 2016. Control of Autophagy in Chlamydomonas Is Mediated through Redox-Dependent Inactivation of the ATG4 Protease. Plant Physiol 172(4), 2219–2234.

Perez-Perez, M.E., Zaffagnini, M., Marchand, C.H., Crespo, J.L., Lemaire, S.D. 2014. The yeast autophagy protease Atg4 is regulated by thioredoxin. Autophagy 10(11), 1953–1964.

Qi, H., Lei, X., Wang, Y., Yu, S., Liu, T., Zhou, S.K., Chen, J.Y., Chen, Q.F., Qiu, R.L., Jiang, L., Xiao, S. 2022. 14-3-3 proteins contribute to autophagy by modulating SINAT-mediated degradation of ATG13. Plant Cell 34(12), 4857–4876.

Qi, H., Xia, F.N., Xie, L.J., Yu, L.J., Chen, Q.F., Zhuang, X.H., Wang, Q., Li, F., Jiang, L., Xie, Q., Xiao, S. 2017. TRAF Family Proteins Regulate Autophagy Dynamics by Modulating AUTOPHAGY PROTEIN6 Stability in Arabidopsis. Plant Cell 29(4), 890–911.

Qi, W., Sun, W. 2025. Analysis of the threshold range of ROS concentration in winter rapeseed of the Brassica napus type. Front Plant Sci 16, 1673768.

Scherz-Shouval, R., Shvets, E., Fass, E., Shorer, H., Gil, L., Elazar, Z. 2007. Reactive oxygen species are essential for autophagy and specifically regulate the activity of Atg4. EMBO J 26(7), 1749–1760.

Song, R.F., Li, T.T., Liu, W.C. 2021. Jasmonic Acid Impairs Arabidopsis Seedling Salt Stress Tolerance Through MYC2-Mediated Repression of CAT2 Expression. Front Plant Sci 12, 730228.

Suttangkakul, A., Li, F., Chung, T., Vierstra, R.D. 2011. The ATG1/ATG13 protein kinase complex is both a regulator and a target of autophagic recycling in Arabidopsis. Plant Cell 23(10), 3761–3779.

Tan, W., Chen, J., Yue, X., Chai, S., Liu, W., Li, C., Yang, F., Gao, Y., Gutierrez Rodriguez, L., Resco de Dios, V., Zhang, D., Yao, Y. 2023. The heat response regulators HSFA1s promote Arabidopsis thermomorphogenesis via stabilizing PIF4 during the day. Sci Adv 9(44), eadh1738.

Wada, S., Hayashida, Y., Izumi, M., Kurusu, T., Hanamata, S., Kanno, K., Kojima, S., Yamaya, T., Kuchitsu, K., Makino, A., Ishida, H. 2015. Autophagy supports biomass production and nitrogen use efficiency at the vegetative stage in rice. Plant Physiol 168(1), 60–73.

Walter, M., Chaban, C., Schutze, K., Batistic, O., Weckermann, K., Nake, C., Blazevic, D., Grefen, C., Schumacher, K., Oecking, C., Harter, K., Kudla, J. 2004. Visualization of protein interactions in living plant cells using bimolecular fluorescence complementation. Plant J 40(3), 428–438.

Wang, P., Liu, W.C., Han, C., Wang, S., Bai, M.Y., Song, C.P. 2024. Reactive oxygen species: Multidimensional regulators of plant adaptation to abiotic stress and development. J Integr Plant Biol 66(3), 330–367.

Wang, Q., Hou, S. 2022. The emerging roles of ATG1/ATG13 kinase complex in plants. J Plant Physiol 271, 153653.

Wang, Q., Qin, Q., Su, M., Li, N., Zhang, J., Liu, Y., Yan, L., Hou, S. 2022. Type one protein phosphatase regulates fixed-carbon starvation-induced autophagy in Arabidopsis. Plant Cell 34(11), 4531–4553.

Woo, J., Park, E., Dinesh-Kumar, S.P. 2014. Differential processing of Arabidopsis ubiquitin-like Atg8 autophagy proteins by Atg4 cysteine proteases. Proc Natl Acad Sci U S A 111(2), 863–868.

Xiong, Y., Contento, A.L., Bassham, D.C. 2007. Disruption of autophagy results in constitutive oxidative stress in Arabidopsis. Autophagy 3(3), 257–258.

Yang, C., Luo, M., Zhuang, X., Li, F., Gao, C. 2020. Transcriptional and Epigenetic Regulation of Autophagy in Plants. Trends Genet 36(9), 676–688.

Yang, H., Zhang, Y., Lyu, S., Mao, Y., Yu, F., Liu, S., Fang, Y., Deng, S. 2025. Arabidopsis CIRP1 E3 ligase modulates drought and oxidative stress tolerance and reactive oxygen species homeostasis by directly degrading catalases. J Integr Plant Biol 67(5), 1274–1289.

Yoshimoto, K., Jikumaru, Y., Kamiya, Y., Kusano, M., Consonni, C., Panstruga, R., Ohsumi, Y., Shirasu, K. 2009. Autophagy negatively regulates cell death by controlling NPR1-dependent salicylic acid signaling during senescence and the innate immune response in Arabidopsis. Plant Cell 21(9), 2914–2927.

Yoshimoto, K., Ohsumi, Y. 2018. Unveiling the molecular mechanisms of plant autophagy – from autophagosomes to vacuoles in plants. Plant and Cell Physiology 59(7), 1337–1344.

Yoshimoto, K., Shibata, M., Kondo, M., Oikawa, K., Sato, M., Toyooka, K., Shirasu, K., Nishimura, M., Ohsumi, Y. 2014. Organ-specific quality control of plant peroxisomes is mediated by autophagy. J Cell Sci 127(Pt 6), 1161–1168.

Yuan, H.M., Liu, W.C., Lu, Y.T. 2017. CATALASE2 Coordinates SA-Mediated Repression of Both Auxin Accumulation and JA Biosynthesis in Plant Defenses. Cell Host Microbe 21(2), 143–155.

Zeng, X., Zeng, Z., Liu, C., Yuan, W., Hou, N., Bian, H., Zhu, M., Han, N. 2017. A barley homolog of yeast ATG6 is involved in multiple abiotic stress responses and stress resistance regulation. Plant Physiol Biochem 115, 97–106.

Zhang, C., Li, S.-Q., Jing, P., Wu, R.-X., Ma, Y.-Q., Wu, J.-X., Song, R.-F., Liu, W.-C. 2026. Autophagy, ROS, and their interplay in plant adaptive responses. Journal of Plant Physiology 316.

Zhang, Y., Song, R.-F., Hu, X.-Y., Zhou, H., Wang, W., Zhang, J., Zhang, X., Li, L.-M., Wang, S.-T., Song, Y., Xiang, F., Xing, J., Long, Y., Zhang, C., Botella, J.R., An, G., Guo, S., Liu, W.-C., Song, C.-P. 2025. Arabidopsis HSFA1b functions as a heat sensor inhibiting OST1-mediated stomatal closure through its adenylate-cyclase activity. Molecular Plant 18(9), 1549–1566.

Zhang, Y., Song, R.F., Yuan, H.M., Li, T.T., Wang, L.F., Lu, K.K., Guo, J.X., Liu, W.C. 2021. Overexpressing the N-terminus of CATALASE2 enhances plant jasmonic acid biosynthesis and resistance to necrotrophic pathogen Botrytis cinerea B05.10. Mol Plant Pathol 22(10), 1226–1238.

