## Supplemental Figures for "Catalase 2-dependent regulation of autophagy in response to carbon starvation in Arabidopsis"

### Supplementary Figure Legends

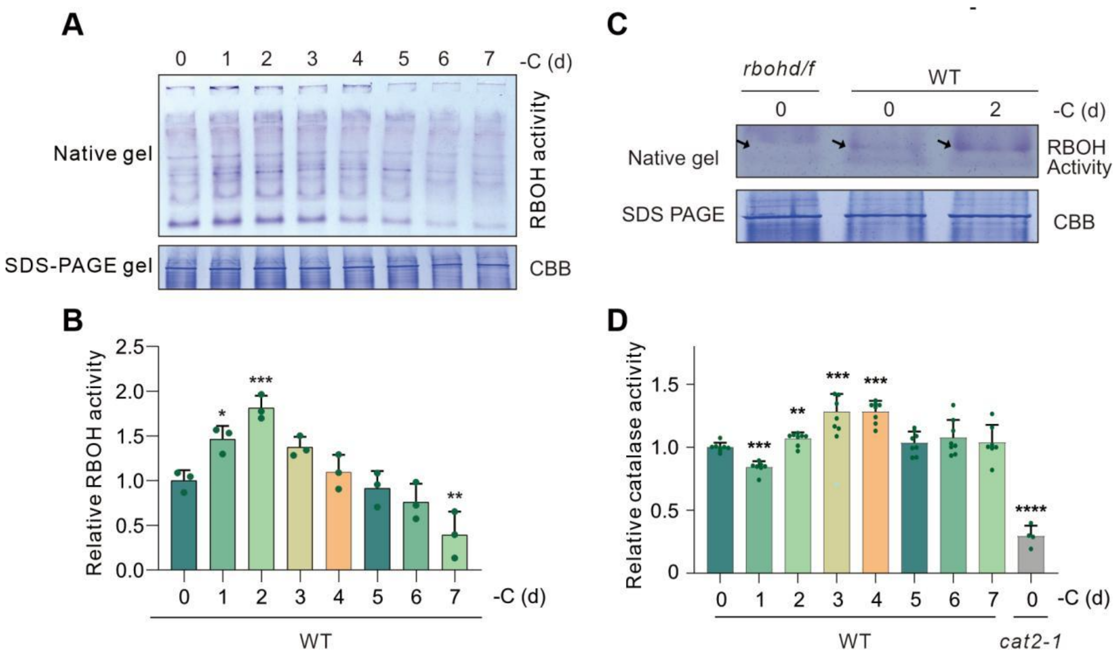


**Figure S1. Analysis of RBOH and catalase activities in plants treated with carbon starvation.**

(**A**) In-gel native assay (top) for RBOH activity in WT seedlings treated with carbon starvation from 0 to 7 d, and the corresponding SDS–PAGE coomassie-blue staining (CBB) (bottom).

(**B**) The relative RBOH activity shown in (A) was determined. Data are means ± SD (n = 3). Bars with different letters indicate significant differences at $p < 0.05$, revealed using Student's t test analysis of variance with a Tukey's multiple comparison test.

(**C**) Representative native gel showing the RBOH activity in the WT and *rbohd/f* mutant plants. Black arrows indicate the absence of specific bands or RBOH activity in the *rbohd/f* mutant compared with the WT.

(**D**) Catalase activity of the WT and *cat2-1* mutant plants treated with carbon starvation from 0 to 7 d. Data are means ± SD (n = 3). Bars with different letters indicate significant differences at $p < 0.05$, revealed using Student's t test analysis of variance with a Tukey's multiple comparison test.

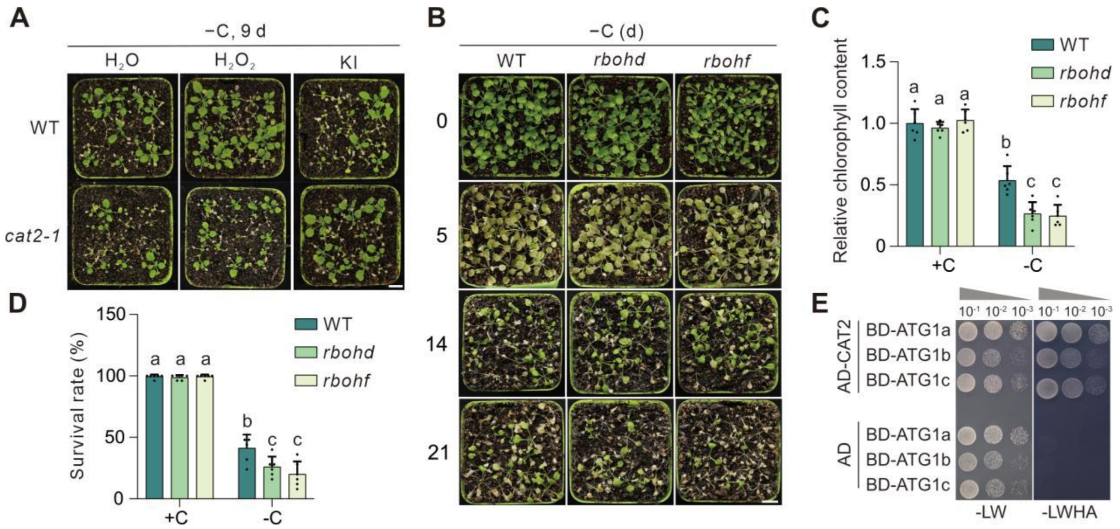


**Figure S2. Carbon starvation phenotypic analysis of the *cat2, rbohd* and *rbohf* mutants.**

(**A**) Representative images showing WT and *cat2* seedlings treated with carbon starvation in the presence or absence of $H_2O_2$ or KI for 9 d. Bar = 1 cm.

(**B**) Time-course analysis of WT, *rbohd* and *rbohf* mutant plants treated with carbon starvation for 0, 5, 14 and 21 d. Bar = 1 cm.

(**C**) The relative chlorophyll content of the WT, *rbohd* and *rbohf* mutant plants treated with carbon starvation for 21 d. Data are mean ± SD (n = 6). Bars with different letters indicate significant differences at $P < 0.05$, revealed using a one-way analysis of variance with a Tukey's multiple comparison test..

(**D**) Survival rates of the WT, *rbohd* and *rbohf* mutant plants treated with carbon starvation for 21 d. Data are mean ± SD (n = 6). Bars with different letters indicate significant differences at $P < 0.05$, revealed using a one-way analysis of variance with a Tukey's multiple comparison test..

(**E**) Yeast two-hybrid assay showing the interaction of CAT2 with ATG1a, ATG1b and ATG1c.

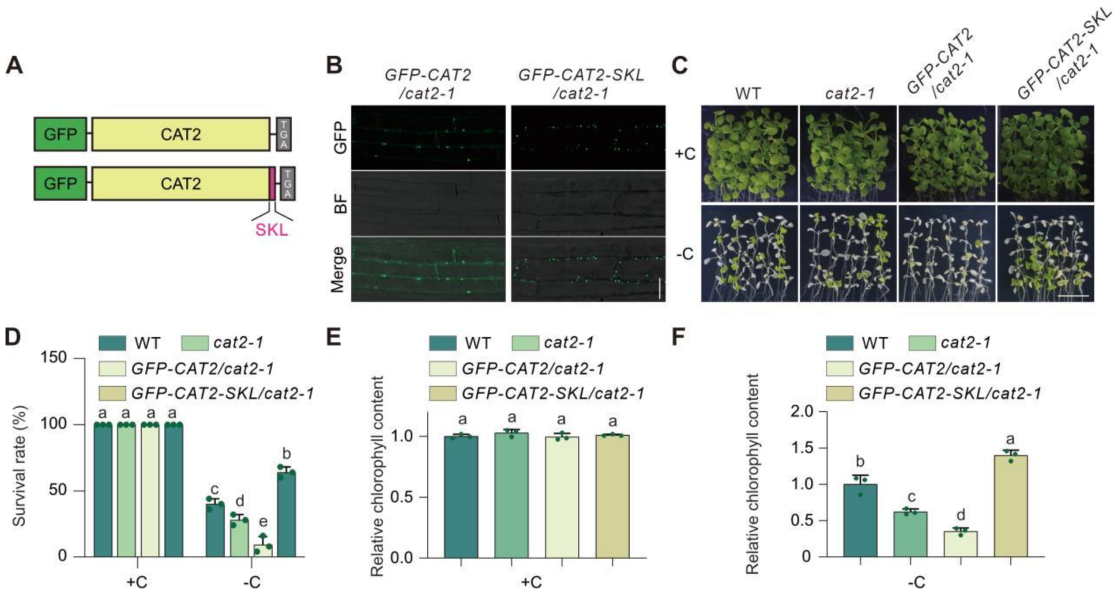


**Figure S3. Expression of peroxisome-localized CAT2 enhances carbon starvation tolerance of the *cat2* mutant.**

(**A**) Schematic diagrams showing the GFP–CAT2 and GFP–CAT2–SKL constructs.

(**B**) Confocal images of the *GFP-CAT2/cat2* and *GFP-CAT2-SKL/cat2* transgenic plant seedlings, showing that GFP-CAT2 has both peroxisomal and cytosolic localizations while GFP–CAT2–SKL is specifically localized in peroxisomes. Bar = 20 μm.

(**C**) Phenotypes of WT, *cat2*, *GFP-CAT2/cat2* and *GFP-CAT2–SKL/cat2* plant seedlings under +C and −C conditions. Bar = 10 mm.

(**D**) Survival rates of WT, *cat2*, *GFP-CAT2/cat2* and *GFP-CAT2–SKL/cat2* plant seedlings treated with or without carbon starvation.under +C and −C conditions. Data are mean ± SD (n = 3). Bars with different letters indicate significant differences at P < 0.05, revealed using a one-way analysis of variance with a Tukey's multiple comparison test.

(**E–F**) Relative chlorophyll content of WT, *cat2*, *GFP-CAT2/cat2* and *GFP-CAT2–SKL/cat2* plant seedlings under +C (E) and −C (F). Data are mean ± SD (n = 3). Bars with different letters indicate significant differences at P < 0.05, revealed using a one-way analysis of variance with a Tukey's multiple comparison test.

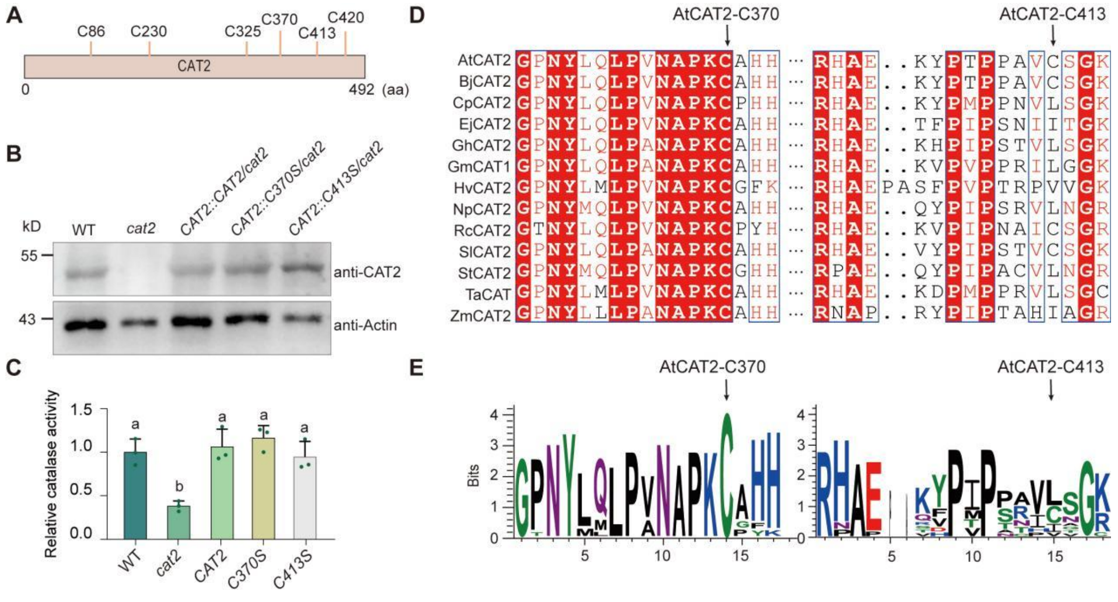


**Figure S4. Analysis of CAT2 protein expression and catalase activity in the transgenic complement lines of the *cat2* mutant.**

(**A**) A schematic diagram showing the positions of six cysteines in CAT2 protein.

(**B**) Immunoblot analysis showing the CAT2 protein abundance in WT, cat2, *CAT2/cat2*, *CAT2-C370S/cat2*, and *CAT2-C413S/cat2* lines using anti-CAT2 antibody. Actin was used as a loading control..

(**C**) Catalase activity in WT, *cat2*, *CAT2/cat2*, *CAT2-C370S/cat2*, and *CAT2-C413S/cat2* lines. Data are mean ± SD (n = 3). Bars with different letters indicate significant differences at $P < 0.05$, revealed using a one-way analysis of variance with a Tukey's multiple comparison test.

(**D–E**) Sequence alignment of CAT2 proteins from multiple plant species, with black arrows indicating the position corresponding to Cys-370 and Cys-413 in Arabidopsis CAT2. At, *Arabidopsis thaliana*; Bj, *Brassica juncea*; Cp, *Cucurbita pepo*; Ej, *Eriobotrya japonica*; Gh, *Gossypium hirsutum*; Gm, *Glycine max*; Hv, *Hordeum vulgare*; Np, *Nicotiana plumbaginifolia*; Rc, *Ricinus communis*; Sl, *Solanum lycopersicum*; St, *Solanum tuberosum*; Ta, *Triticum aestivum*; Zm, *Zea mays.*
